# Cytokinin N-conjugate Form Activity, Metabolism, and Signaling During Leaf Senescence

**DOI:** 10.64898/2026.05.08.723873

**Authors:** Omar Hasannin, Ivan Petřík, Miroslav Strnad, Ondřej Novák, Martin Černý, Aaron M. Rashotte

**Affiliations:** 101 Rouse Life Sciences, Department of Biological Sciences, Auburn University, Auburn, AL 36849 USA; Department of Molecular Biology and Radiobiology, Faculty of AgriSciences, Mendel University in Brno, Zemedelska 1, CZ-61300, Brno, Czech Republic; Laboratory of Growth Regulators, Faculty of Science, Palacký University, Šlechtitelů 27, CZ-77900, Olomouc, Czech Republic; Laboratory of Growth Regulators, Institute of Experimental Botany, The Czech Academy of Sciences, Šlechtitelů 27, CZ-77900, Olomouc, Czech Republic

**Author notes:** Corresponding Author: Aaron M. Rashotte 101 Rouse Life Sciences, Department of Biological Sciences, Auburn University, USA. Email: Omar Hasannin Ivan Petřík Miroslav Strnad Ondřej Novák Martin Černý Aaron M. Rashotte.

**Keywords:** Cytokinin, Cytokinin N-conjugates, Metabolome, Transcriptome, Proteome, Senescence

## Abstract

Cytokinin (CK) N-glucosides are the most abundant CK metabolites in *Arabidopsis* and most angiosperms, yet their role in cytokinin activity and response is unclear. Here, we examined metabolomic, transcriptomic, and proteomic profiles of seven CK N-glucoside conjugates in detached *Arabidopsis* leaves across a 144-hour dark-induced senescence (DIS) timecourse. All tested N-glucosides were found to undergo a slow conversion to their corresponding base forms at position-dependent rates, with N9-glucosides releasing base faster than their corresponding N7-glucosides. Conversion during DIS was strictly isoform-specific and not accompanied by coordinated induction of CK biosynthesis genes, arguing against *de novo* synthesis as the source of accumulated base. Despite progressive base accumulation, N-glucoside-treated leaves produced substantially fewer Differentially Expressed Genes than direct base application at comparable base concentrations, revealing a disconnect between hormone presence and transcriptional output. Unbiased model comparison identified the base:glucoside ratio as a stronger predictor of CK-Two Component Signaling (TCS) gene expression than absolute base concentration, though modulated by base-type-specific receptor affinities. Early proteomic profiling further revealed a coordinated response shared across N-glucosides but largely absent from base treatments. Together, these findings support that CK N-glucosides as kinetically slow, position-dependent reservoirs whose presence in abundance modulate activation of CK-TCS elicited by bioactive forms.

**Highlights:** Physiology, metabolomic, transcriptomic, and proteomic findings here support CK N-glucosides as kinetically slow, position-dependent reservoirs whose presence in abundance modulate activation of CK-TCS elicited by bioactive forms.

## Introduction

Cytokinins (CKs) are adenine-derived plant hormones with central roles in cell division, shoot and root development, leaf senescence, and stress responses (Werner and Schmülling, 2009; Kieber and Schaller, 2018). Bioactive forms in Arabidopsis are the free bases *trans*-zeatin (*t*Z), N6-isopentenyladenine (iP), dihydrozeatin (DHZ) and, to a lesser extent, *cis*-zeatin (*c*Z) (Hasannin et al., 2026) that are perceived by Arabidopsis Histidine Kinase (AHKs) receptors and signal through a multistep two-component system to activate the type-A and type-B response regulators (ARRs) that drive most CK-responsive transcriptional output (Hwang et al., 2012). CK bases are subject to metabolic homeostasis through irreversible side-chain cleavage by Cytokinin Oxidase/Dehydrogenase (CKX) family, ribosylation, O-glucosylation, and N-glucosylation at positions N7 or N9 on the adenine moiety, catalyzed primarily by UGT76C1 and UGT76C2 in Arabidopsis (Hou et al., 2004; Takei et al., 2004; Wang et al., 2011; Wang et al., 2013).

Despite the large investment made in producing N-glucosides, typically composing the largest pool of CK metabolites in Arabidopsis and most angiosperms (Novák et al., 2008; Skalický et al., 2023); the biological function of N-glucosides has remained poorly understood. Early biochemical work treated them as irreversible inactivation products of base forms (Letham et al., 1983). However, recent studies reported bioactivity of exogenously treated N-glucosides, albeit sometimes weak or tissue– and species-dependent (Hallmark et al., 2020; Pokorná et al., 2020; Khanna et al., 2025). It is unclear how observed functional activity of N-glucoside is achieved since no N-glucosides have affinity to the AHK receptors (Spíchal et al., 2004; Šmehilová et al., 2016). The irreversibility view has, however, been challenged by recent reports that tZ-type N-glucosides can be hydrolyzed back to free *t*Z, with N9-glucosides hydrolyzed faster than N7-glucosides on minute-to-hour timescales (Hošek et al., 2020; Hoyerová and Hošek, 2020). Whether such reverse conversion generalizes across the iP, DHZ and *c*Z families as well as the consequences for transcriptional signaling *in planta* have not been examined.

In this study, we ask three connected questions about CK N-glucosides during dark-induced senescence, where these forms have shown the most activity to delay senescence. First, do all N-glucosides undergo conversion to their corresponding bases under physiological conditions, and on what timescale? Second, do N-glucosides themselves elicit any cellular response that is distinct from the response to their corresponding free base? Third, why do plants invest resources in synthesizing and maintaining large pools of N-glucosides? Using detached *Arabidopsis* rosette leaves in a classical Dark-Induced Senescence (DIS) assay, treatment with each of seven N-glucosides across a 144-hour time course was paired with metabolomic, transcriptomic and proteomic analyses. From this, we show that N-glucosides function as kinetically slow, position-dependent reservoirs of bioactive CKs and that transcriptional output of the released base is temporally and quantitatively decoupled from its concentration.

## Methods and Materials

### Dark-induced leaf senescence (DIS) assays and plant growth conditions

Wild-type Arabidopsis seeds (Col-0) were planted in plastic trays with Pro-Mix BX and placed in growth chambers under a 16/8h light cycle (100 μmol m^−2^ s^−1^) with temperatures of 22°C/18°C. Dark-induced leaf senescence (DIS) bioassays were modified for use with Arabidopsis as previously described in (Fletcher and McCullagh, 1971; Johnston et al., 2026). Briefly, 4-week-old leaves were detached and floated on 3 mL of 3 mM MES buffer solution (pH 5.7) in 6-well culture plates, one leaf/well; only leaf positions 5 and 6 were used unless otherwise indicated. Cytokinin treatments (*t*Z7G, *t*Z9G, iP7G, iP9G, DHZ7G, DHZ9G, and *c*Z9G) were applied individually to a final concentration of 1 µM and compared to DMSO or Ethanol (0.1%) as a vehicle control and placed under growth chamber conditions in the dark. Measurements of DIS were performed at 2, 48, 96, or 144 hours after treatment/dark incubation, as noted below. All CKs were obtained from OlChemIm (Olomouc, Czech Republic) as analytical standards with >95 % purity and with N-glucosides free from base form contamination. Adenine was obtained from AmBeed Co. with >99% purity.

#### Photosystem II efficiency (Fv/Fm), chlorophyll content, and ion leakage measurements

In a dark room, FluorCam FC 1000-H was used to measure Fv/Fm; leaf areas were selected manually, and the scaling bar was set to 100. Chlorophyll (Chl) was extracted according to (Nayek et al., 2014; Johnston et al., 2026). Briefly, leaf weights were recorded at the start of the experiment. Leaves were harvested at each of the designated time points, followed by Chl extraction using methanol incubation overnight. Debris was pelleted by centrifugation at maximum speed for 10 min, and supernatant was used to measure total Chl normalized to fresh weight The number of biological replicates is indicated in the figure legend of each experiment; generally, at least four independent biological replicates were performed per treatment/time-point. ANOVA followed by post-hoc test was used to determine statistical differences.

#### Quantification of cytokinin profiles

Whole leaf samples (20-30 mg fresh weight) were ground in liquid nitrogen then lyophilized and stored until quantification was performed using three biological replicates (n=18 independent leaves per replicate). Lyophilized tissues were extracted in 0.5mL extraction solvent consisting of 5% formic acid in 75% methanol (both v/v). Four zirconium oxide homogenization beads and a mixture of stable isotope-labelled internal standards of CKs were added to each sample (0.2 pmol CK bases, ribosides and N-glucosides, 0.5 pmol CK O-glucosides and nucleotides). The standards were obtained from Olchemim Ltd. (Olomouc, Czech Republic) and from the Faculty of Science, Palacký University in Olomouc (Czech Republic) with distinct compounds as listed in (Svačinová et al. 2012). The samples were extracted using mix-mode solid phase extraction (Dobrev and Kamínek 2002). First, the samples were shaken in Retsch MM 400 oscillation bead mill (Retsch, Haan, Germany) for 5 min at 27 Hz, 4°C, sonicated for 3 min and incubated for 30 min at 4°C. Then the extracts were centrifuged at 20,000 rpm, 4°C for 15 min (Allegra 64R benchtop centrifuge, Beckman Coulter, USA). The supernatant was diluted with 2.5 mL 1 M aqueous formic acid and loaded onto activated Oasis® MCX 30 mg/1 cc extraction cartridge (Waters, Milford, USA). The activation was performed using 1 mL methanol, and 2mL 1 M aqueous formic acid. After the loading of the sample, the cartridge was washed with 1 mL 1M aqueous formic acid and 1 mL 80% methanol (v/v). The CKs were eluted with addition of 1mL aqueous 0.35 M ammonia and 2 mL 0.35 M ammonia in 60% methanol (v/v). The eluate was evaporated to dryness using SpeedVac concentrator (RC1010 Centrivap Jouan, ThermoFisher, USA) and reconstituted in 40 µl of 10% aqueous methanol (v/v). The sample was transferred into LC vial equipped with a glass insert (Chromservis Ltd., Czech Republic) and then analyzed using ultra-high performance liquid chromatography Acquity I-Class system (Waters, Milford, USA) coupled with Xevo TQ-S series tandem mass spectrometer (Waters, Manchester, UK). The chromatographic conditions and mass spectrometry settings were set as previously published (Svačinová et al. 2012). Data were acquired and processed in multiple reaction monitoring using MassLynx V4.2 software. The final concentration of plant hormones (pmol/g of fresh weight) was calculated using isotope dilution method (Rittenberg and Foster 1940). Values were log-transformed, and non-detected values were replaced with the detection limits to enable statistical comparisons determined by ANOVA followed by a post-hoc test.

#### RNA extraction and RNA-seq transcriptome analysis

RNA was isolated from frozen whole leaf powder (approximately 60 mg) of three biological replicates, n=18 leaves per 1 biological replicate; three independent biological replicates were extracted using RNeasy Plant Mini Kits (Qiagen) according to the manufacturer’s instructions. RNA was sent to Novogene, Inc for Illumina sequencing.

Raw RNA-sequencing reads were processed by Novogene, including raw gene counts, differential expression of genes between treatments, and GO enrichment. Briefly, high-quality clean transcript reads for all downstream analyses were obtained by first removing sequencing adapters, reads containing poly-N, and low-quality reads. Using HiSAT2 *v2.0.5* (Pertea et al. 2016), the *Arabidopsis thaliana* TAIR10 reference genome was indexed and paired-end clean reads were aligned. The mapped reads were assembled by StringTie *v1.3.3b* in a reference-based approach and counted using featureCounts *v1.5.0-p3* (Liao, Smyth, and Shi 2014; Pertea et al. 2015). Differential expression analysis of two conditions/groups (three biological replicates per condition) was performed using the DESeq2 R package (1.20.0) (Love, Huber, and Anders 2014). The resulting P-values were adjusted using the Benjamini and Hochberg’s approach for controlling the false discovery rate. Genes with an adjusted P-value (padj) ≤0.05 found by DESeq2 were assigned as differentially expressed (Robinson, McCarthy, and Smyth 2010). The P values were adjusted using the Benjamini & Hochberg method and a corrected P-value of 0.05 and absolute fold change of 2 were set as the threshold for significantly differential expression.

#### Proteome analysis

The extraction of proteins from lyophilized tissues was conducted according to the protocol delineated by (Berková et al., 2023). In brief, tissue samples underwent homogenization, and aliquots ranging between 10-20 mg were lyophilized. Subsequently, these were extracted utilizing a mixture of tert-butyl methyl ether and methanol (3:1 ratio). The precipitated protein pellets were then resolubilized in a solution composed of 8 M urea, 10 mM dithiothreitol (DTT), and 100 mM ammonium bicarbonate. Following solubilization, the proteins underwent alkylation and were digested with trypsin. Each sample, containing precisely 5 µg of peptides, was subjected to analysis via nanoflow reverse-phase liquid chromatography coupled with mass spectrometry (LC-MS/MS). The chromatographic separation was achieved using a 15 cm C18 Zorbax column on an Agilent system, interfaced with a Dionex Ultimate 3000 RSLC nano-UPLC and an Orbitrap Fusion Lumos Tribrid Mass Spectrometer, equipped with a FAIMS Pro Interface (Thermo Fisher Scientific, Waltham, MA, USA). The analysis utilized alternating FAIMS compensation voltages of −40, −50, and −75 V. The acquired MS/MS spectra were subsequently recalibrated and interrogated against the Araport 11 protein database, alongside a database of common contaminants, using Proteome Discoverer 2.5. Algorithms such as Sequest HT, MS Amanda 2.0, and MSFragger facilitated the search (Dorfer et al., 2014; Kong et al., 2017). Quantitative analysis was restricted to proteins identified by at least two unique peptides The comprehensive mass spectrometry proteomics dataset has been made accessible through the ProteomeXchange Consortium via the PRIDE partner repository, under the dataset identifier PXD053034. Differential Enriched Protein analysis was conducted using DEP package in R.

#### Statistical modeling analysis

Apparent conversion rate constants (*k*_conv_) were estimated under a steady-state assumption in which continuous uptake from the application medium maintains the tissue N-glucoside pool while slow first-order hydrolysis generates free base. Under this assumption, *k*_conv_ = (linear rate of base accumulation above control) / (mean tissue N-glucoside concentration across the time course). Linear rates were estimated by ordinary least-squares regression of replicate-level excess base on time. Conversion half-life is reported as ln(2) / *k*_conv_.

In mass balance analysis, for each treatment and timepoint, the percentage of the initial N-glucoside pool required to account for all observed excess base under a closed fixed-pool model was calculated as (excess base above control) / (initial excess N-glucoside above control at the 2-hour timepoint) × 100. The coefficient of variation (CV) of N-glucoside measurements at each timepoint was calculated as (standard deviation / mean) × 100. A treatment is considered consistent with closed-pool conversion only if the percentage of pool required to account for the observed base remains below the measurement CV; treatments exceeding 100% of the initial pool require continuous-uptake replenishment and were interpreted under a steady-state framework.

Statistical modeling of CK-TCS gene expression was performed in R (version 4.4.0) using the lme4 package for mixed-effects model fitting and the lmerTest package for Satterthwaite-approximated p-values. Marginal and conditional R² were calculated with the MuMIn package following (Nakagawa et al., 2017).

To avoid selection bias, we used a pre-defined set of canonical CK-responsive genes from the meta-analysis of (Bhargava et al., 2013); the strict-tier set comprised 10 genes responsive to CK in ≥13 of the meta-analyzed datasets (seven type-A ARRs, CKX4, ASL9, and AT4G03610). For each treatment × timepoint combination, free base and total N-glucoside concentrations were calculated from LC-MS measurements of the corresponding base family (sum of N7– and N9-glucoside isomers). Gene expression was expressed as the z-score difference between treatment and time-matched control, with each gene treated as an independent observation. The base-to-N-glucoside ratio was calculated as base concentration divided by total N-glucoside concentration plus 0.01 to avoid division by zero, then log – transformed. All concentration-based predictors were z-score standardized to allow comparison of standardized effect sizes.

Mixed-effects models took the general form gene_z_score ∼ fixed_predictors + tp_factor + (1 | gene), with timepoint as a fixed factor and gene identity as a random intercept. Cross-family pooled models additionally included (1 | treatment) as a random effect; within-family models omitted this term due to the small number of treatments per family. Model comparisons were done via AIC, BIC, and likelihood ratio tests, using the convention that ΔAIC < 4 indicates substantial support for the inferior model and ΔAIC > 10 indicates essentially no support. To identify the strongest predictor of CK-TCS gene expression, seven candidate model specifications were compared: null (timepoint only), base only, N-glucoside only, ratio only, additive base + N-glucoside, base × N-glucoside interaction, and base + ratio. To directly compare ratio versus base concentration as predictor while controlling for family identity, models of identical complexity differing only in the within-family quantitative predictor (Base + Family vs Ratio + Family; Base × Family vs Ratio × Family) were compared via AIC.

To quantify systematic deviation of N-glucoside-treated samples from base-only predictions, an ordinary least-squares regression model (gene_z_score ∼ base_concentration + factor(timepoint) + factor(gene)) was fit to base-treatment data within each family and used to predict expression for both base and N-glucoside treatments. Residuals were compared between treatment classes using both Wilcoxon rank-sum and two-sample t-tests. Details of all models are in Supplemental File 5

#### Protoplast isolation and transient expression

The protocol was adapted from (Yoo et al., 2007; Panda et al., 2024). Briefly, ∼30 3-weeks old Arabidopsis leaves were cut horizontally into 0.5-1mm strips, excluding the leaf tip and base. Leaf strips submerged in enzyme solution (1.5% Cellulase (Onozuka R-10), 0.4% macerozyme R-10, 0.4M Mannitol, 20mM KCl, 10mM CalC2, and 0.1% BSA) were vacuumed in the dark with gentle shaking (40-50 rpm) for at least 3 hours. The digested protoplasts were filtered through a 40 µm cell strainer and washed with W5 buffer (2 mM MES, 154 mM NaCl, 125 mM CaCl2, and 5 mM KCl) by centrifugation at 100 × g for 7 minutes. Next, protoplasts were resuspended in 6 mL of 0.55 M freshly prepared sucrose and centrifuged at 100 × g for 30 minutes. Healthy protoplasts were extracted from the interface, followed by a W5 wash, and counted using a hemacytometer. 1 × 106/mL protoplasts were resuspended in MMG solution (4 mM MES, 0.4 M mannitol, and 15 mM MgCl2). 110 µL PEG 40% with 0.2 M mannitol, and 100 mM CaCl2, were added to 100 µL of protoplasts and 10 µL of DNA (15 µg TCSn::LUC and 5 µg UBQ10::GUS), followed by dark incubation for 20 minutes, and quenching with 0.9 mL W5 solution. Protoplasts were incubated in WI solution (4 mM MES, 0.5 M mannitol, and 20 mM KCl) supplemented with the indicated concentrations of tZ and/or tZ7G. For each treatment, a total of 5 µL of compound mix was added to 1 mL of protoplast suspension to ensure equal solvent volumes across all conditions. Details of stock concentrations, volumes, and mixing schemes are provided in **Supplementary File 5**. After 6 hours of dark incubation, protoplasts were harvested for LUC (Promega Cat # E1500) and GUS measurement as described in (Yoo et al., 2007). Five biological replicates per condition from 5 different isolations were conducted.

To evaluate transfection efficiency, we transfected 10 µg of 35S::GFP and incubated overnight. 45%-60% of cells were successfully transfected. UBQ10::GUS and 35S::GFP were obtained from ABRC (CD3-1038 and CD3-911, respectively). TCSn::LUC was obtained from (Zürcher et al., 2013).

#### Quantitative realtime PCR

Treated detached leaves (positions 5-6) were incubated in the dark for 2 hours with the indicated treatment. At least 2 leaves per treatment/condition were ground together and counted as one biological replicate (repeated 3 times). Followed by RNA extraction as previously described. 500 ng of RNA was used for cDNA synthesis using Quantbio qScript cDNA Synthesis Kit. 5 µL of 1:10 dilution of cDNA was used for qPCR mix, using PerfeCTa SYBR® Green FastMix. All primers used in the study are in **(Supplemental File 5).** TUB4 served as an internal control; relative expression was calculated using ΔΔCt method. Statistical analysis was conducted by comparing ΔCt values using ANOVA post-hoc.

## Results

### Cytokinin N-glucosides generally show weak and inconsistent anti-senescence activity

To evaluate the physiological activity of CK N-glucosides at the whole-leaf level, we performed (DIS) assays, a well-established assay to evaluate CK activity (Gan and Amasino, 1995). For this, detached *Arabidopsis* leaves (positions 5 and 6) were treated with 1 µM of each of seven N-glucoside conjugates (*t*Z7G, *t*Z9G, iP7G, iP9G, DHZ7G, DHZ9G, and *c*Z9G) and effects were compared to their corresponding base treatments and an ethanol negative control using F_v_/F_m_, and chlorophyll content, as senescence-progression markers **(Figure 1).** The active base forms *t*Z, iP, and DHZ produced strong and consistent protection against senescence (Hasannin et al., 2026). In contrast, most N-glucosides failed to significantly preserve chlorophyll content or F_v_/F_m_ levels relative to the negative control. Only *t*Z7G and *t*Z9G produced significant differences in F_v_/F_m_ against the negative control in specific time points, and these effects were inconsistent across leaf developmental stages **(Supplemental Fig. S1 and S2).**

**Figure 1.**
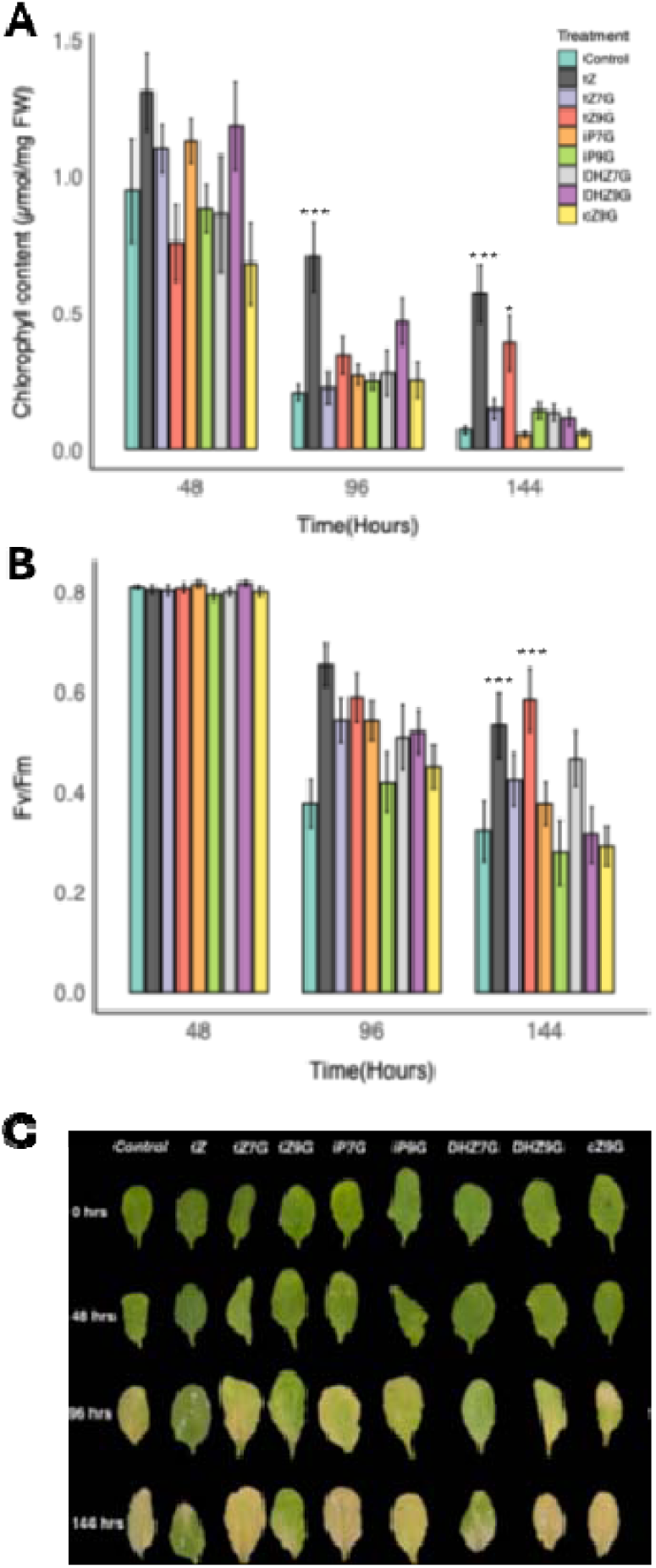
N-glucosides show inconsistent and weak activity in delaying leaf senescence. (**A-C**) Quantification of senescence progression across the 144-hour time course. Total chlorophyll content (µmol/mg FW) **(A),** Photosystem II efficiency (Fv/Fm) **(B)** of detached Arabidopsis leaves (positions 5-6) treated with 1 µM tZ, tZ7G, tZ9G, iP7G, iP9G, DHZ7G, DHZ9G, cZ9G, or negative control (NC) and incubated in the dark for 48, 96, and 144 hours. **(C)** Representative images of detached leaves from 4-week-old Arabidopsis plants treated with 1 µM of the indicated cytokinin bases form treatment across the time-course of dark-induced senescence (DIS). Data represent the mean ± SE of at least 4 biological replicates. Asterisks indicate statistical significance compared to the negative control (NC) as determined by one-way ANOVA followed by Bonferroni post-hoc test for (A-C) (*P < 0.05, **P< 0.001, ***P < 0.0001).

### N-glucoside pools are stable while their corresponding bases accumulate gradually over senescence

To investigate the metabolic dynamics of exogenously applied N-glucosides, we quantified endogenous CK profiles at the same times across 144-hour DIS assay as examined physiology for physiology in Figure 1 and in other omics analysis **(Supplemental File S2).** All N-glucoside treatments elevated the corresponding endogenous measured conjugate levels in treated leaves above control, confirming uptake and tissue accumulation of the applied compound **(Fig. 2A).**

**Figure 2.**
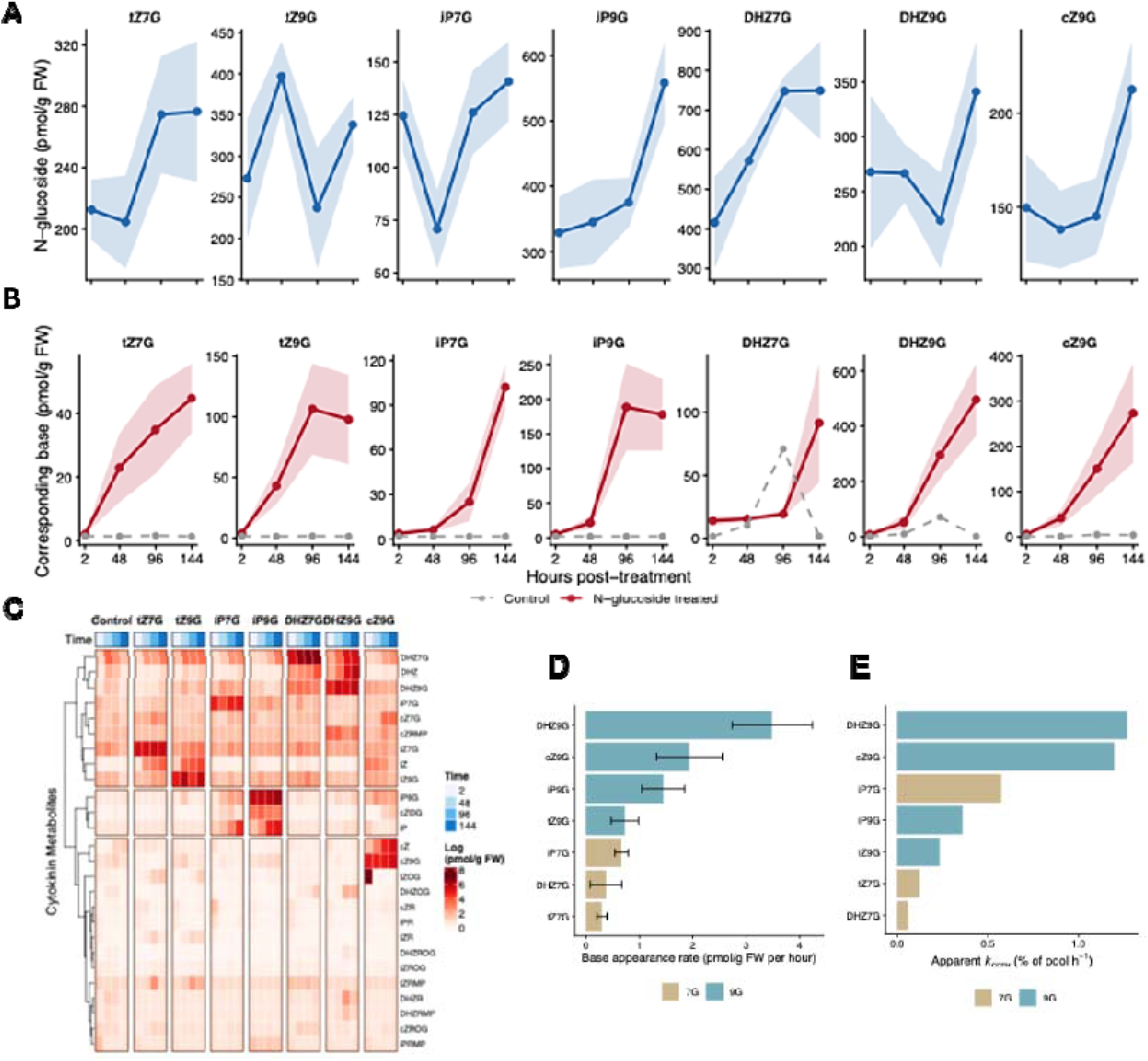
N-glucoside metabolism: pool dynamics, base release, and conversion kinetics. **(A)** Time course of N-glucoside pool concentration (pmol/g fresh weight) for each of the seven N-glucoside treatments across 2, 48, 96, and 144 hours. Values are means ± SD (n = 3). Linear regression statistics for pool stability are reported in Table S1. **(B)** Time course of corresponding free base accumulation (pmol/g FW) in N-glucoside-treated leaves. Each panel shows accumulation of the parent base (e.g., tZ in tZ7G– and tZ9G-treated samples) across the 144-hour time course. **(C)** Heatmap of endogenous cytokinin metabolite levels (log-transformed) across treatments at 2, 48, 96, and 144 hours post-treatment. **(D)** Mean base appearance rates (pmol/g FW/h) computed by linear regression across the 144-hour time course for each N-glucoside treatment. Bars are colored by glucoside position (N7 vs N9). Error bars represent SE of the regression slope. N9-glucosides show consistently higher base appearance rates than corresponding N7-glucosides for tZ and iP families; DHZ shows the largest N9/N7 ratio (∼9-fold). **(E)** Apparent *k*_conv_ indicating the percentage of converted N-glucosides into their corresponding base forms under a steady replenishment assumption.

Accumulation of N-glucoside conjugates differed substantially across compounds and developed over the DIS time course **(Supplemental Table S1).** At 2 h post-treatment, mean pool concentrations ranged from a low of 125±38 pmol/g FW for iP7G to a high of 416±259 pmol/g FW for DHZ7G, with only marginal between-treatment differences (one-way ANOVA, p = 0.053). By 48 h, between-treatment differences became highly significant (p = 3.0 × 10□□) and remained so through 144 h (p = 1.1 × 10□□). DHZ7G accumulated to consistently higher concentrations than all other N-glucosides, reaching 748±86 pmol/g FW at 96 h, while iP7G and cZ9G accumulated to the lowest concentrations across all timepoints (≤ 213 pmol/g FW). N9-glucosides of *t*Z and iP accumulated to higher tissue concentrations than the corresponding N7-glucosides at later timepoints, while DHZ7G exceeded DHZ9G across all timepoints.

Unlike base treatments, where the applied free base declined steadily in leaf samples across DIS from 2 to 144 hours consistent with active CKX-mediated turnover (Hasannin et al., 2026), N-glucoside pools showed no significant decline for any treatment over DIS (linear regression on raw concentrations, *p* > 0.05 for slope < 0 in all seven treatments; **Supplemental Table S1 and Supplemental Table S2).** Two N-glucosides (iP9G, *p* = 0.011 and DHZ7G, *p* = 0.008) actually showed significant net accumulation across the time course, increasing by 70% to 80% relative to the 2-hour level **(Supplemental Table S2).** Pool stability or net accumulation in the face of cumulative substrate flux is consistent with the known resistance of N-glucosides to CKX-mediated degradation (Galuszka et al., 2007) combined with continuous uptake from the medium.

Despite this pool stability, all seven N-glucoside treatments resulted in a gradual and steady increase of their corresponding base form across the 144-hour time course **(Fig. 2B).** The accumulation was strictly isoform-specific: *t*Z7G and *t*Z9G elevated *t*Z, iP7G and iP9G elevated iP, DHZ7G and DHZ9G elevated DHZ, and *c*Z9G elevated *c*Z, with no substantial cross-base accumulation. Across all seven N-glucoside treatments and four timepoints, only one significant elevation of a non-corresponding base was detected, an increase of *t*Z in *c*Z9G-treated samples at 48 hours (compared to control, FDR-adjusted *p* = 0.020; Fig. 2C). The strict isoform-specificity of base accumulation argues against generalized activation of the IPT/LOG biosynthetic pathway to produce more base, which would be expected to elevate multiple CK types simultaneously (Sakakibara, 2021).

To quantify the dynamics of base appearance, we calculated the rate of base accumulation above control for each N-glucoside treatment by linear regression of replicate-level excess base on time **(Fig. 2D).** Rates differed substantially across treatments, ranging from 0.30±0.09 pmol g^−1^ FW h^−1^ for tZ7G to 3.48±0.75 pmol g^−1^ FW h^−1^ for DHZ9G. Overall two consistent patterns emerged. First, a Timepoint×base type interaction term in a pooled linear model, which was significant (*p* = 0.024) confirming that the rate of base accumulation depends on the parent base. Second, N9-glucosides released back to their corresponding base at substantially faster rates than the corresponding N7-glucosides for all three families where both positions were tested (Fig 2D). DHZ9G was 9.3-fold faster than DHZ7G, tZ9G was 2.5-fold faster than *t*Z7G, and iP9G was 2.2-fold faster than iP7G. This positional difference supports the previous biochemical observations of t*Z*-N-glucosides over short timescales (Hošek et al., 2020) to the full N-glucoside family across a 144-hour window in planta. It additionally indicates that the N9-glycosidic bond is consistently more labile to *in vivo* hydrolysis than the N7 bond regardless of the base it carries.

### Base accumulation is consistent with slow conversion under steady-state uptake and is not attributable to *de novo* biosynthesis

A natural question is whether the base appearing in N-glucoside-treated leaves arises from direct hydrolysis of the applied substrate or from *de novo* CK biosynthesis induced by the treatment. Under a simple closed-system model in which the applied N-glucoside is the only source of base, the percentage of the initial glucoside pool required to account for all observed excess base by 144 hours ranged from 21.8% (*t*Z7G) to 188% (*c*Z9G), with five of seven treatments exceeding 50% and three exceeding 100% **(Supplemental Table S3).** Several treatments therefore require more substrate to be consumed than was what is present at the start of the experiment, which is impossible under a strictly closed-pool assumption. Combined with the absence of pool decline, this rules out fixed-pool conversion.

In our experimental design, however, leaves remain in contact with the applied hormones in the medium throughout the 144-hour time course, allowing continuous uptake of N-glucoside from the medium into the tissue. We therefore considered a steady-state model in which uptake from the medium replenishes whatever substrate is lost to slow hydrolysis. This would leave the tissue pool approximately constant (or, where uptake exceeds hydrolysis, slowly accumulating, as observed for iP9G and DHZ7G) while base is generated continuously and accumulates because its production rate exceeds its CKX-mediated clearance. Under this assumption, we estimated the apparent conversion rate constant (*k*_conv_) for each treatment as the ratio of the linear base appearance rate to the mean N-glucoside concentration during the time course. The estimated *k*_conv_ values ranged from 0.06% pool h^−1^ for DHZ7G (half-life ∼1146 h) to 1.27% pool h^−1^ for DHZ9G (half-life ∼55 h) **(Fig. 2E and Supplemental Table S4).** At these rates, only a few percent of the pool convert within any single inter-timepoint interval, well within the measurement noise of the glucoside pool itself (CV typically 20-60%), while cumulative conversion across 144 hours, sustained by continuous uptake, can quantitatively account for the observed base accumulation without contribution from *de novo* synthesis.

To test whether *de novo* biosynthesis nonetheless contributes to base accumulation, we examined the expression of known CK biosynthesis genes across all timepoints in N-glucoside-treated samples via RNAseq transcriptome analysis of leaf DIS treated samples **(Figure 3).** Since base accumulation in N-glucoside treated samples was observed as early as 2 hours, if de novo biosynthesis is occurring, it is expected that N-glucosides treatment would trigger the induction of CK-biosynthesis genes. However, we found no evidence of coordinated upward induction of the biosynthetic pathway **(Fig. 3A)**. Mean expression of the biosynthesis-gene set was indistinguishable from control at every timepoint and in every N-glucoside treatment **(Supplemental Fig. S3).** Individual genes did show scattered changes, but these were bidirectional, inconsistent across treatments, and similar in scale and direction to feedback responses observed in base-treated samples at matched timepoints. As such they likely reflect canonical CK-induced negative feedback on biosynthesis rather than activation. Taken together with the strict isoform-specificity of base accumulation, as absence of coordinated biosynthesis activation makes novo synthesis less likely as the source of the accumulated base and supports a slow hydrolysis of the applied N-glucoside under steady-state uptake system.

**Figure 3.**
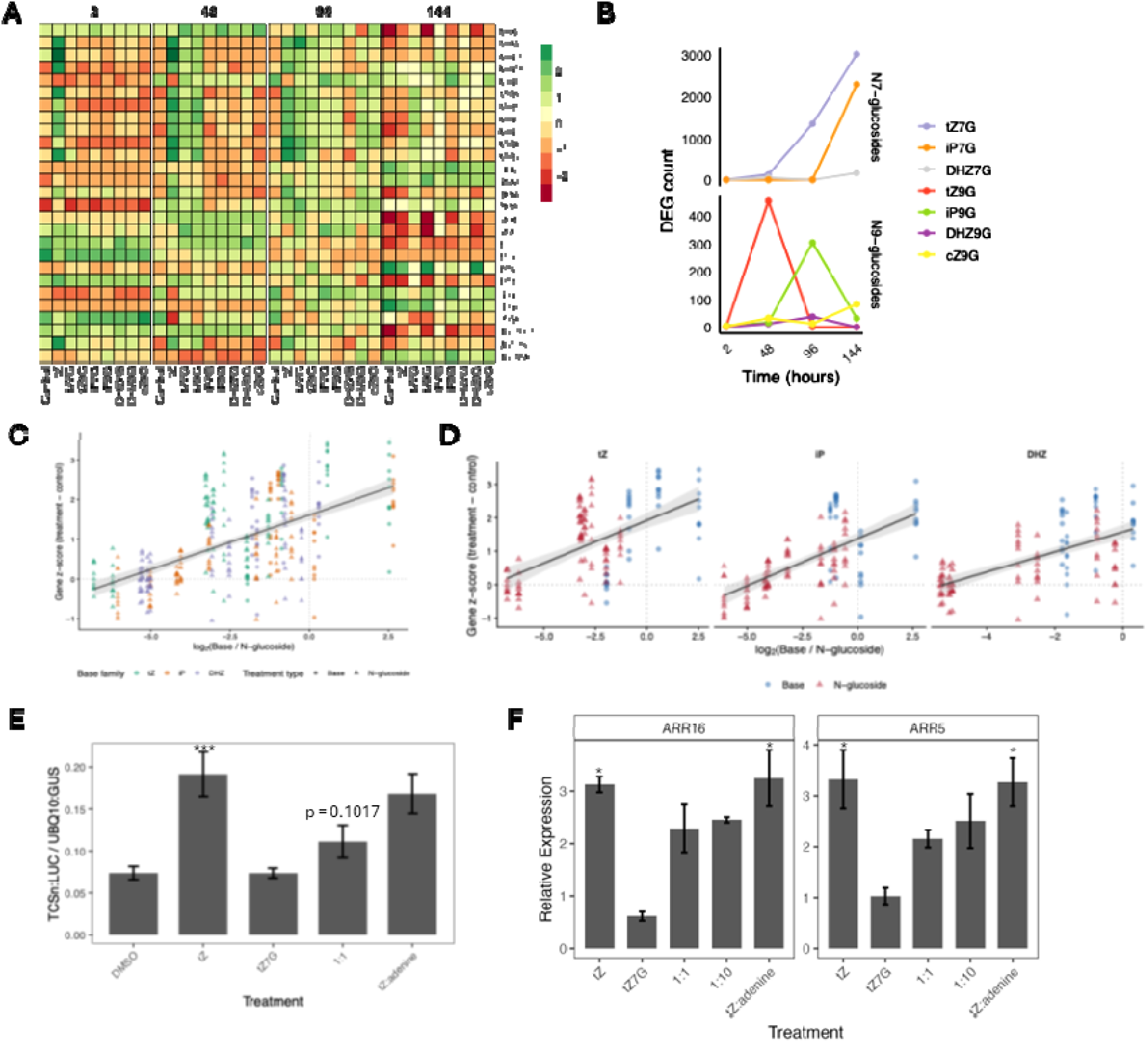
Cytokinin gene expression by N-glucosides and the disconnect between bulk-tissue base and transcriptional output. **(A)** Heatmap of canonical cytokinin signaling and homeostasis gene expression (Type-A ARRs, IPTs, LOGs, CKXs, UGTs, and Type-B ARRs) across all treatments at 2, 48, 96, and 144 hours. Color scale: z-score difference from time-matched control. **(B)** Number of differentially expressed genes (DEGs; |log2 fold-change| > 1 and adjusted p < 0.05) per treatment per timepoint. N-glucosides show delayed and inconsistent induction of DEGs. tZ7G at 144 hours produces an exception of 3,024 DEGs. **(C)** Mean z-score of a pre-defined CK-responsive gene set (Bhargava et al. 2013; ≥13 microarray datasets responding to cytokinin; n = 10 genes); showing the effect of ratio Log2 base/N-glucoside in a scatter plot vs. CK-TCS gene z-score (treatment − time-matched control), faceted by base family (DHZ, iP, tZ) **(D),** Separate linear regressions are shown for base treatments (blue circles) and N glucoside treatments (red triangles). N-glucoside-treated samples consistently fall below the base-only regression line. **(E)** Co-application of equal concentrations of tZ and tZ7G to protoplasts reduces TCSn signal compared to tZ alone. **(F)** Expression of RR16 and RR5 is reduced when co-applying tZ and tZ7G compared to tZ alone, while co-application of equal concentrations of tZ and adenine did not affect regulation levels. Data represent the mean ± SE for 6 biological replicates (6 hours after treatment) **(D)** and 3 biological replicates (two hours after treatment) **(E).** Asterisks indicate statistical significance compared to the NC as determined by Student t-test in (*P < 0.05, **P< 0.001, ***P < 0.0001).

In contrast to CK biosynthesis genes, CK degradation and conjugation genes showed a clear and reproducible temporal pattern in N-glucoside-treated leaves **(Fig. 3A)**. *CKX4* was induced above control at 96 and 144 hours in nearly every N-glucoside treatment, with treatment-specific late inductions of additional CKX family members. Both N-glucosylation enzymes *UGT76C1* and *UGT76C2* were also strongly induced at 96 and 144 hours, with the largest single response observed in iP7G-treated leaves (*UGT76C1* z-score difference of 3.03 at 144 h; **Fig 3A and Supplemental Fig. S4).** Because CKX and UGT76C2 genes are themselves canonical CK-responsive targets (Bhargava et al., 2013), their late induction provides indirect evidence that the conversion-derived base eventually reaches concentrations sufficient to engage homeostatic clearance. The concurrent induction of *UGT76C1*/*C2* also raises the possibility of a partial re-conjugation cycle in which freshly released base is recaptured into the N-glucoside pool, contributing to the observed pool stability.

### Transcriptional responses to N-glucosides are temporally disconnected from base accumulation

Given that N-glucoside-derived base reach concentrations sufficient to activate CKX and UGT homeostatic genes, we asked whether the broader transcriptional response tracks base accumulation in a manner predictable from base-treatment kinetics. We compared the total number of differentially expressed genes (DEGs; padj < 0.05, |log_2_FC| > 1) and the expression of canonical CK-TCS marker genes between base and N-glucoside treatments across the 144-hour time course **(Fig.3B).**

Base treatments produced an expected temporal pattern: few DEGs at 2 hours, a strong response peak at 48 and 96 hours (*t*Z: 995 and 1853 DEGs; iP: 1047 and 1326; DHZ: 1044 and 1205, respectively), and a sharp decline by 144 hours (*t*Z: 7 DEGs; DHZ: 26; iP: 243**; Supplemental Fig S5,** Hasannin et al., 2026). This pattern reflects the time required for signaling cascade propagation followed by clearance of the applied base by CKX, with peak transcriptional output occurring well after the 2-hour base concentration maximum. The weak *c*Z produced minimal DEGs across all timepoints (1-75), consistent with its low affinity for AHK receptors (Spíchal et al., 2004; Lomin et al., 2015).

N-glucoside treatments, by contrast, produced fragmented and temporally inconsistent transcriptional responses **(Fig. 3B).** Most conjugates produced few or no DEGs at early timepoints despite measurable base accumulation: iP7G generated zero DEGs at 2 and 48 hours, DHZ9G zero at 2 hours, c*Z*9G only 2 and 32 DEGs at 2 and 48 hours respectively. Several treatments produced no detectable DEGs at timepoints when the corresponding base had reached substantial concentrations. t*Z*9G generated 452 DEGs at 48 hours when tissue t*Z* levels were approximately 43 pmol g^−1^ FW, but zero DEGs at both 96 and 144 hours when tZ levels were comparable or higher **(Fig. 3B and Fig 2).** iP7G remained transcriptionally silent through 48 hours despite gradual iP accumulation, then produced a burst of 2,298 DEGs at 144 hours when iP exceeded 100 pmol g^−1^ FW. Additionally, the global transcriptional expression profiles of bioactive base forms (*t*Z, iP, and DHZ) differed substantially in each time point compared to the expression profiles of N-glucosides **(Supplemental Fig. S6).** The sporadic response pattern contrasts sharply with the predictable transcriptional arc of base treatments and indicates that base concentration is not, by itself, a sufficient predictor of transcriptional output when base is delivered through N-glucoside conversion.

Plotting base concentration against DEG count made the disconnect explicit **(Supplemental Fig. S7).** For base treatments, the DEG count was uncorrelated with base concentration (Spearman ρ = –0.10, *p* = 0.70, n = 16), reflecting a temporal lag between peak concentration (early) and peak transcriptional output (48-96 h). In contrast for N-glucoside treatments, the corresponding base concentration and DEG count were positively correlated (ρ = +0.58, *p* = 0.0045, n = 22). However, at any given base concentration, most N-glucoside treatments produced substantially fewer DEGs than the corresponding base treatment. For example, DHZ9G treatment at 96 hours contained 295 pmol g^−1^ FW of DHZ but produced only 37 DEGs, whereas DHZ treatment at 48 hours contained a comparable 397 pmol g^−1^ FW and produced 1,044 DEGs **(Supplemental Fig. S7).** Similarly, iP9G treatment at 96 hours had 189 pmol g^−1^ FW of iP and produced 303 DEGs, compared with 1,047 DEGs from iP base treatment with comparable iP levels at 48 hours. The exception was *t*Z7G, which by 144 hours produced 3,024 DEGs; a magnitude exceeding any base-treatment timepoint at a *t*Z concentration of only 44.7 pmol g^−1^ FW. In addition, iP7G also produced high number of DEGs with a low base concentration.

Examining canonical CK-TCS marker expression directly produced the same picture with a cleaner signal **(Supplemental Fig. S8).** The mean z-score difference from control across *ARR5, ARR6, ARR7, ARR15, ARR16, AHK4, CKX4* and *UGT76C2* expression was consistently elevated in base treatments (z-score difference ranging from 1.0 to 2.8 above control across active timepoints) and uncorrelated with base concentration (Spearman ρ = +0.18, *p* = 0.51). In N-glucoside treatments, TCS marker induction was generally lower; most treatments at 2 hours showed mean z-score differences near zero or slightly negative, indicating no early activation of canonical CK signaling. TCS expression in N-glucoside treatments rose gradually over the time course in parallel with base accumulation (Spearman ρ = +0.68, *p* = 0.0001, n = 22), but in most cases remained below the levels achieved by direct base treatments at comparable base concentrations. tZ7G again was the exception, reaching base-like TCS marker induction by 96 and144 hours (z-score differences of 2.0 and 2.4, respectively).

### Late N-glucoside transcriptional responses are consistent with delayed canonical CK signaling

To test whether the late, large transcriptional responses observed in *t*Z7G and iP7G represent activation of a glucoside-specific signaling pathway distinct from canonical CK signaling, or, alternatively, delayed activation of the canonical pathway by released base, we characterized the DEG set shared between *t*Z7G and iP7G at 144 hours that was absent from any DEGs from base treatment at any timepoint. Of the 3,024 DEGs in *t*Z7G at 144 hours and 2,298 DEGs in iP7G at 144 hours, 982 genes were shared between them and not present in any of the four base treatments at any timepoint **(Supplemental Fig. S9).** Gene Ontology analysis revealed that these genes are enriched for Autophagy, and stress-associated terms **(Supplemental Fig. S10).** If these genes represented a distinct glucoside-specific signaling program, they would be expected to appear early in N-glucoside treatments and to enrich for regulators outside the canonical CK-TCS circuit. Instead, almost all of these genes (982 of 982 for iP7G; 982 for *t*Z7G with 234 also detected at 96 h and 6 at 48 h) appeared exclusively at the latest timepoint **(Fig 3B).** The direction of regulation was approximately balanced (553 up, 429 down for both treatments), and no canonical CK-TCS genes were present in this set. This suggests that whatever CK-related signaling is occurring at 144 hours is being routed through the same canonical pathway activated by direct base application, rather than through a parallel glucoside-specific mechanism.

### Base-to-N-glucoside ratio is the strongest predictor of CK-TCS gene expression

The temporal-disconnect between base accumulation and transcriptional output suggested that base concentration alone is insufficient to predict CK signaling. To test this quantitatively, we fit gene-level mixed-effects models to a pre-defined set of canonical CK-responsive genes from an independent meta-analysis (Bhargava et al., 2013; genes responsive to CK in ≥13 of the meta-analyzed datasets, n = 10 genes including seven type-A ARRs, CKX4, ASL9, and AT4G03610). Models compared whether base concentration alone, N-glucoside concentration alone, the base-to-N-glucoside ratio, additive base+N-glucoside terms, or a base×N-glucoside interaction best predicted gene z-score differences from time-matched control, with random effects on gene identity.

Across all eleven treatments and four timepoints (n = 440 observations), the base-to-N-glucoside ratio decisively outperformed base concentration as a predictor of CK-TCS gene expression. Direct comparison of mixed-effects models showed that a Ratio+Family model outperformed an equivalent Base+Family model by ΔAIC = 42, with marginal R² increasing from 0.42 to 0.47. In a joint model containing both predictors, the standardized ratio coefficient (β = 0.45, t = 7.22, p < 10^-^¹²) was 2.6-fold larger than the standardized base coefficient (β = 0.17, t = 2.64, p = 0.009), and adding ratio to a base-only model improved fit by χ² = 49.2 (p = 2.3 × 10^-^¹²) while adding base to a ratio-only model improved fit only modestly (χ² = 6.9, p = 0.009, **Supplemental Table S5).** Plotting CK-TCS gene z-score against log (base/N-glucoside) across all observations revealed a unified positive relationship **(Fig. 3C and Supplemental Table S6)**, confirming that the ratio of base to N-glucoside captures the dominant quantitative relationship between CK metabolites and transcriptional output.

The pooled relationship was modulated by base family identity in a manner consistent with known CK receptor affinities **(Supplemental Tables S7 and S8).** Per-family regression slopes were similar for iP (0.32) and DHZ (0.33), shallower for *t*Z (0.22), and shallowest for *c*Z (0.16, **Supplemental Table S9),** with family-specific intercepts following previously reported potency hierarchy of CK bases for AHK receptors (*t*Z > iP ≈ DHZ > *c*Z; (Spíchal et al., 2004; Lomin et al., 2015). At equivalent log_□_ ratios, tZ-family treatments produced systematically higher transcriptional responses than iP, DHZ, or *c*Z family treatments **(Fig. 3D).** Adding family identity as a fixed effect to a ratio-only model significantly improved fit (ΔAIC = 104, χ² = 110, p < 10^-1^□) and adding family-specific slopes provided a further improvement (ΔAIC = 15, p = 1.3 × 10^-^□). The full description of the relationship is therefore ratio-driven with family-dependent intercepts that reflect intrinsic differences in CK receptor affinities.

Within each bioactive family, N-glucoside-treated samples produced significantly lower transcriptional response than predicted by their base concentration alone. Residuals from a base-only regression model were significantly negative in every bioactive family (Wilcoxon test against base residuals; *t*Z: residual = –0.09, p = 0.047; iP: residual = –0.57, p = 1.1 × 10^-^□; DHZ: residual = –0.75, p = 1.2 × 10^-^□; **Supplemental Fig. S11).**

To test the effect of base: glucoside ratio directly, we performed two complementary co-application experiments. In Arabidopsis mesophyll protoplasts transformed with the synthetic CK reporter pTCSn::LUC (Zürcher et al., 2013). Following 6 hours of application of 100 nM *t*Z produced robust reporter activation (∼2.5-fold over DMSO control), while application of 100 nM *t*Z7G alone produced no detectable activation, confirming that *t*Z7G is not directly bioactive on the CK signaling pathway. Co-application of *t*Z and *t*Z7G (1:1) reduced pTCSn::LUC activity by 41% relative to *t*Z alone, while co-application of *t*Z with an equal concentration of adenine did not reduce reporter activity, ruling out non-specific effects **(Fig. 3E).**

To verify this finding in planta, we measured transcript levels of two canonical type-A ARR genes (ARR5 and ARR16) by qRT-PCR in detached Arabidopsis leaves treated with *t*Z alone, *t*Z7G alone, mixed *t*Z:*t*Z7G at 1:1 and 1:10 molar ratios, or *t*Z with equal adenine concentration, 2 hours following treatment. Both ARR5 and ARR16 were strongly induced by *t*Z (∼3.0-fold) and not significantly induced by *t*Z7G alone (≤ 1-fold). The 1:1 and 1:10 *t*Z:*t*Z7G mixtures reduced induction of ARR5 by 36% and 24% and ARR16 by 27% and 22% respectively, while the *t*Z + adenine control showed no significant difference compared to *t*Z alone **(Fig. 3F).** Together, these findings provide a direct validation of the omics-based finding that the relative quantity of base versus N-glucoside, not base concentration alone, determines transcriptional output.

### N-glucosides elicit unique and consistent induction of proteins after 2 hours of treatment

Given that N-glucoside-treated leaves produced few or no transcriptional responses at 2 hours despite demonstrable uptake, we asked whether earlier signaling events might be detectable at the protein level. We performed quantitative proteomic profiling of the same 2-hour leaf samples used for the transcriptomic and metabolic analyses. 126 proteins responded significantly in terms of altered abundance levels to at least one of the eleven treatments (Figure 4). Per-treatment counts were highly heterogeneous among direct base treatments (*c*Z: 0; iP: 1; *t*Z: 27; DHZ: 52) but more uniform across N-glucoside treatments (range 11 to 35; **Fig. 4B).** N-glucoside responses predominantly increases protein abundance; across the seven N-glucosides, 130 of 158 (82%) significant protein changes were increases above control **(Fig 4A and Supplemental Fig S12),** while base treatments produced more bidirectional patterns (e.g., *t*Z: 18 up / 9 down; DHZ: 35 up / 17 down). Direct iP application produced a single significant protein at 2 hours, whereas iP7G and iP9G produced 22 and 20 respectively; direct *c*Z application produced no differential protein abundances, while *c*Z9G produced 35 **(Fig. 4B).** The substantially larger early proteomic response to N-glucosides than to their corresponding free bases, applied at the same concentration in the same experiment, cannot be straightforwardly attributed to slow base release alone, since the bases themselves produce essentially no DEPs at this timepoint while their conjugates produce coordinated responses **(Fig. 4C and Supplemental Fig. S13).**

**Figure 4.**
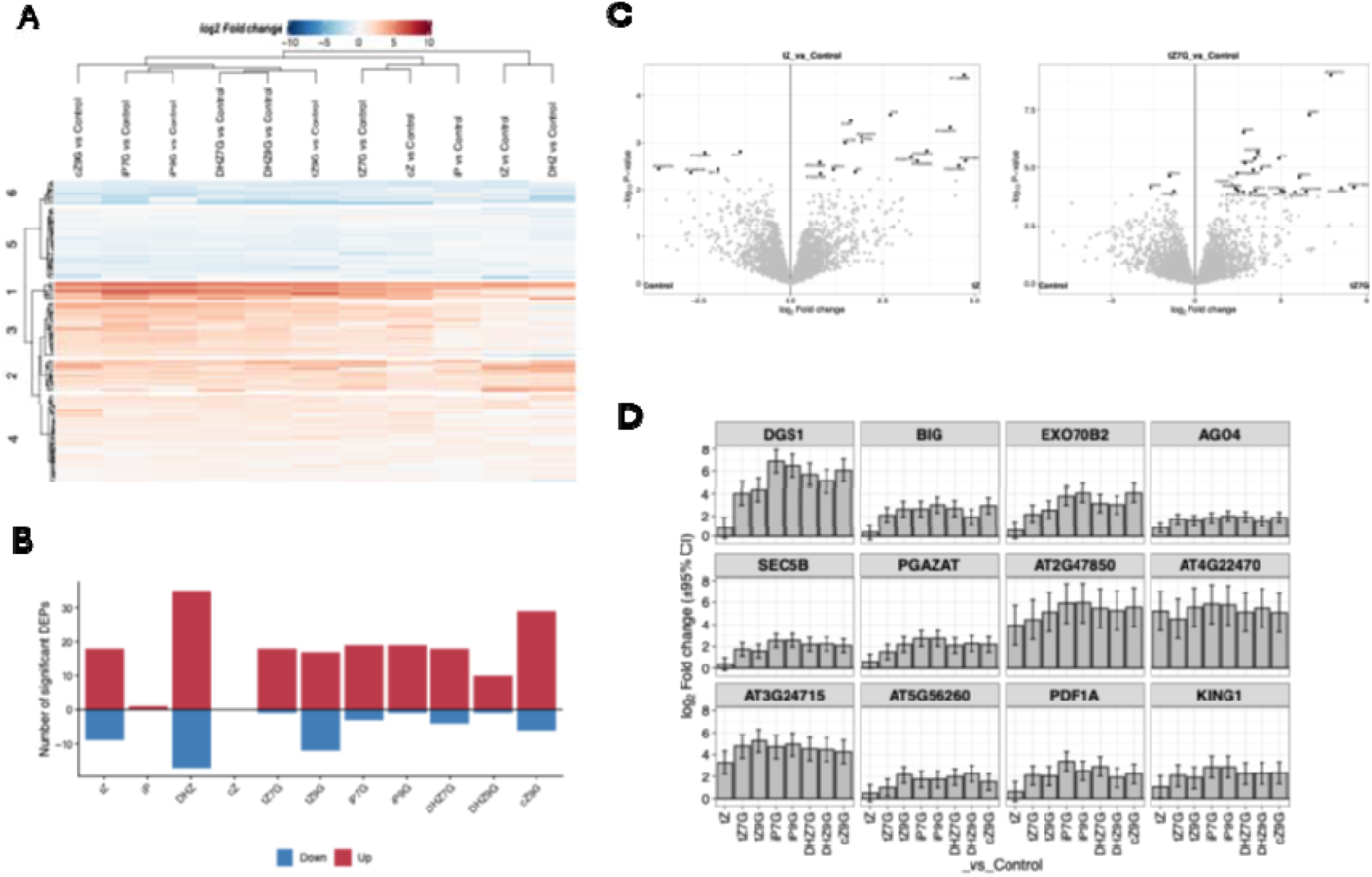
Distinct early proteomic response signature in N-glucosides treated samples. (A) cluster plot of proteomic changes in response to base form and N-glucosides based on log2fold change compared to negative control. N-glucosides form a separate cluster from base forms in their proteomic signature. **(B)** Number of significant protein changes per treatment at 2 hours post-treatment (adjusted p < 0.05, |log2 fold-change| > log2(1.3)), with up– and down-regulated proteins shown in different colors. Bases produce highly heterogeneous numbers (cZ: 0; iP: 1; tZ: 27; DHZ: 52), while all seven N-glucosides produce 11-35 significant proteins, predominantly upregulated. **(C)** volcano plot of significant Differentially Enriched Proteins (DEPs) in response to tZ and tZ7G compared to negative control. **(D)** Bar plots showing mean log2 fold-change (vs. time-matched control) ± 95% confidence interval for the 12 core N-glucoside-induced proteins (significant in ≥5 of 7 N-glucoside treatments) across all 11 treatments. Treatment order: tZ, tZ7G, tZ9G, iP, iP7G, iP9G, DHZ7G, DHZ9G, cZ, cZ9G. The core comprises proteins involved in vesicle trafficking and exocytosis (BIG, EXO70B2, SEC5B), cell wall and membrane organization (PGAZAT), lipid handling (DGS1), chromatin regulation (AGO4), bioenergetic regulation (KING1), antimicrobial response (PDF1A), and four uncharacterized proteins (AT2G47850, AT4G22470, AT3G24715, AT5G56260).

To investigate whether N-glucoside responses converge on a shared protein set, we identified proteins significant in at least five of the seven N-glucoside treatments. Thirteen proteins met this criterion, all thirteen abundances were enriched in every N-glucoside treatment in which they were significant, with mean log2 fold-changes ranging from +1.5 to +5.7 **(Fig. 4D, Supplemental File S4).** Six of these thirteen proteins (AT2G47850, DGS1, AT4G22470, AT3G24715, EXO70B2, SEC5B) were significantly enriched in all seven N-glucoside treatments. The same analysis applied to base treatments identified only one protein (AT3G24715) significant in three or more of the four bases, and no protein significant in all four. The N-glucoside class therefore shows a coordinated, base-identity-independent proteomic response that the corresponding direct base treatments do not produce.

Inspection of the response pattern across all eleven treatments confirms this asymmetry **(Supplemental Fig. S12).** The 13 core N-glucoside-response proteins show consistent enriched abundance across all seven N-glucosides, including N-glucosides whose corresponding direct base produced essentially no early proteomic response (iP7G/iP9G versus iP; *c*Z9G versus *c*Z). For 10 of the 13 core proteins, response in the four direct base treatments was either absent or substantially weaker than in the N-glucosides. The three exceptions (AT3G24715, AT2G47850, AT4G22470) showed partial overlap, with significant enrichment in two or three direct base treatments alongside their consistent N-glucoside response. The set overlap analysis shows the asymmetry at the global level: of the 126 significant affected proteins, 59 responded to at least one N-glucoside but no base treatment, 55 responded to at least one base but no N-glucoside, and 12 responded to both classes.

The 13 core N-glucoside-response proteins span functional categories that do not match the canonical CK-TCS transcriptional signature. They include components of vesicle trafficking and exocytosis (BIG, EXO70B2, SEC5B), cell wall and membrane organization (PGAZAT), lipid handling (DGS1), and chromatin and RNA regulation (AGO4) **(Fig 4D)** and were among the most strongly enriched proteins. None of the canonical CK-responsive type-A ARRs, AHK receptors, CKX or UGT76C transcripts have been previously associated with rapid protein-level induction in response to CK treatment, and none appeared in the core set here (Cerny et al., 2011). The early proteomic response to N-glucoside application is therefore qualitatively distinct from the signature expected of canonical CK base perception.

These observations show that the absence of an early transcriptional response to N-glucosides does not reflect an absence of cellular response. Cells detect N-glucoside application within 2 hours and produce a coordinated, predominantly upregulating proteomic adjustment that is shared across N-glucoside isoforms and is largely absent from direct base treatments. Whether this response reflects perception of N-glucosides themselves through a mechanism distinct from canonical CK base perception, a stress or transport response triggered by uptake of glucoside conjugates, a non-transcriptional consequence of trace base release that the corresponding direct base treatments do not engage at the same timepoint, or some combination of these, cannot be resolved from the present data.

## Discussion

This study provides a systematic in planta characterization of CK N-glucoside conversion dynamics across multiple N-glucoside types and an extended physiological time course, resulting in three principal findings. First that exogenous application of a CK N-glucoside causes a slow, position-dependent accumulation to their corresponding base form with N9 conjugates consistently more labile than N7. Second, that conversion-derived base accumulates gradually without coordinated induction of CK biosynthesis genes, supporting hydrolysis rather than *de novo* synthesis as the source. Finally, that the transcriptional response to this gradually accumulating base is attenuated and temporally disconnected from base concentration relative to direct base application; based on our modeling, the ratio of base to corresponding glucoside level is a better predictor of CK-TCS gene expression than base concentration alone. Together, these results support an interpretation of N-glucosides release or conversion back to their respective base forms as a kinetically slow process, serving as biologically reversible reservoirs that modulate the transcriptional activity of bioactive CKs.

The apparent conversion rate constants we report (0.06-1.27% of pool h^−1^; corresponding half-lives 55-1146 hours, **Fig. 2D**) place CK N-glucoside hydrolysis on a substantially slower timescale than canonical CK signaling, which operates within minutes to hours. Under continuous a N-glucoside supply, free base is generated slowly enough that it is plausibly cleared by CKX or recaptured by UGT76C1/C2 (both of which are induced at late timepoints in our data) before reaching receptor-activating concentrations in the relevant cellular compartment. The position dependence we describe of N9 > N7 across *t*Z, iP, and DHZ families, with fold differences of 2.2–9.3 generalizes an observation previously documented for *t*Z-glucosides over short timescales (Hošek et al., 2020) to the broader N-glucoside family and to a 144-hour experimental window in which physiologically meaningful base accumulation can be measured. Notably, our modeling assumes that there is linear and continuous replenishment of the converted N-glucosides from the application medium, which may be plausible given we added 1 µM of hormones that are theoretically resistant to degradation (Galuszka et al., 2007), yet not necessary true; therefore, conversion percentages should be interpreted with caution.

The absence of coordinated early CK biosynthesis pathway gene induction from N-conjugate application together with strict isoform-specificity of base accumulation places strong constraints on the source of the accumulating base under these conditions. A *de novo* biosynthesis explanation would predict simultaneous elevation of multiple CK types, since IPT enzymes generate iPRMP and tZRMP precursors that are processed by LOG and CYP735A enzymes into multiple downstream isoforms (Zhao et al., 2024). Instead, we observe that each N-glucoside elevates mainly its corresponding base, in a pattern that is incompatible with pathway activation but compatible with direct hydrolysis of the applied substrate. The mass-balance impossibility of fixed-pool conversion (several treatments require >100% of the initial pool to be consumed by 144 hours), combined with the absence of significant pool decline and in two cases significant pool accumulation **(Fig 2),** points specifically to a steady-state system in which continuous uptake from the application medium replenishes the substrate as it is slowly hydrolyzed. Whether the same kinetic system operates in intact plants under endogenous conditions where uptake-from-medium does not apply is an open question, but our results show that under conditions of sustained substrate availability, N-glucosides can supply free base at biologically meaningful rates over physiological timescales.

The original biological motivation for this study was the long-standing puzzle of why N-glucosides are the single most abundant CK metabolites in most plant tissues (Skalický et al., 2023), despite their apparent weak activity in some bioassays and inactivity in others (Pokorná et al., 2020). Our data supports a model where the ratio of free bases to N-glucosides might play an important role in modulating the expression of TCS genes rather than base form concentration alone **(Fig 3).**

Within this framework, the well-documented high abundance of N-glucosides in plant tissues becomes biologically interpretable, such that N-glucosides may serve as both a slow-release reservoir of bioactive CKs and as a buffering pool that constrains the maximum signaling output achievable from any given concentration of bioactive base. Despite that we show co-application has the ability to reduce activity in protoplast and *in planta* only in using a *t*Z:*t*Z7G example. Our regression modeling show that this effect is general across N7– and N9-glucosides and across CK-families **(Fig. 3).**

Although it is less clear how this is achieved specifically since direct receptor antagonism appears unlikely based on receptor binding studies showing that N-glucosides do not displace *t*Z from AHK receptor occupancy (Šmehilová et al., 2016). Yet it is plausible that N-glucosides alter cellular states that would modulate the localization of released base forms. As we have shown, despite near absence of early transcriptional induction, all seven N-glucoside treatments produced a coordinated, predominantly enrichment in the abundance of 13-protein core shared across at least five of the seven N-glucosides are absent from comparable analyses of direct base treatments. 82% of significant N-glucoside-induced protein changes were up-regulations, while base treatments produced more bidirectional patterns and considerable treatment-specific heterogeneity (*c*Z producing zero significant changes, iP producing one, DHZ producing 52). This asymmetry suggests that whatever the mechanism, N-glucosides engage a coordinated cellular program that direct application of their corresponding bases does not, at least at this early timepoint and in this experimental system.

Several of these proteins are involved in vesicle trafficking and transport, prompting us to speculate that N-glucosides may manipulate base localization by sequestering them in organelles lacking AHK receptors, such as the vacuole. This hypothesis cannot be resolved through bulk hormone quantification and instead requires organelle-targeted measurements when free bases and N-glucosides are co-applied. A related possibility is that N-glucosides are hydrolyzed and then release base forms in the vacuole, preventing receptor engagement. This would explain why N-glucosides show their strongest relative activity late in our examined DIS assay, at a final leaf senescence stage where rapid apoptosis ruptures the vacuole (Hara-Nishimura and Hatsugai, 2011; Zhuang and Jiang, 2019), releasing the converted bases. In assays without such membrane disruption like root assays and shoot regeneration, N-glucosides show correspondingly no to weaker activity (Hallmark et al., 2020; Hallmark and Rashotte, 2020; Pokorná et al., 2020).

In summary our data support that N-glucosides are not metabolically inert; they undergo measurable conversion to active bases. Whether slow-release-reservoir activity alone fully explains the magnitude of N-glucoside abundance observed across plant tissues is, however, less clear. The conversion rates we measure are slow enough that such a reservoir would be most useful on developmental rather than acute timescales, buffering long-term hormonal balance, sustaining basal signaling during stress, providing a transportable storage form between tissues, or enabling rapid release under specific physiological conditions that we have not probed here. Discriminating among these specific functional roles will require tissue-resolved, developmentally staged, and stress-responsive analyses that go beyond the application-and-measure approach used in the study.

### Supplemental data

**Supplemental Figure S1.** Activity of N-glucosides-treated leaves in dark-induced senescence assay on leaves 3-4.

**Supplemental Figure S2.** Activity of N-glucosides-treated leaves in dark-induced senescence assay on leaves 7-8.

**Supplemental Figure S3.** Average z-score of cytokinin biosynthesis-associated genes in response to N-glucosides and base form treatment.

**Supplemental Figure S4.** Expression of cytokinin degradation genes across time in response to N-glucoside and base form treatments.

**Supplemental Figure S5.** Number of Differentially Expressed Genes in response to base forms.

**Supplemental Figure S6.** Block partial least square analysis comparing the global expression profiles of base form and N-glucoside treatments.

**Supplemental Figure S7.** Plotting number of Differentially Expressed Genes versus accumulated base forms in response to exogenous base and N-glucoside treatments.

**Supplemental Figure S8.** Plotting mean expression of Cytokinin Two Component Signaling Genes versus accumulated base forms in response to exogenous base and N-glucoside treatments.

**Supplemental Figure S9.** Upset plot showing the shared and unique numbers of Differentially Expressed Genes between tZ7G, iP7G and bioactive base forms.

**Supplemental Figure S10.** Gene Ontology analysis on unique genes co-regulated by tZ7G and iP7G but not by bioactive base forms.

**Supplemental Figure S11.** Residuals of linear regression modeling.

**Supplemental Figure S12.** Heat map showing the differential protein abundance in N-glucosides and base form treated-samples.

**Supplemental Figure S13.** Volcano plots showing the differentially abundant proteins in response to N-glucoside-treated samples.

**Supplemental Table S1.** Tissue accumulation of N-glucoside conjugates across the time course.

**Supplemental Table S2.** Pool stability of N-glucosides across the 144-hour time course.

**Supplemental Table S3.** Mass balance of N-glucoside conversion to free base over 144 hours.

**Supplemental Table S4.** Conversion kinetics of N-glucosides to free bases under detached-leaf dark-induced senescence conditions.

**Supplemental Table S5.** Coefficients of the joint mixed-effects model relating standardized base concentration, standardized log (base/N-glucoside) ratio, and base family identity to CK-TCS gene z-score expression.

**Supplemental Table S6.** Comparison of mixed-effects models predicting CK-TCS gene z-score from base concentration, base-to-N-glucoside ratio, and base family identity.

**Supplemental Table S7.** Comparison of ratio-only versus ratio + family versus ratio × family mixed-effects models.

**Supplemental Table S8.** Likelihood ratio test comparing ratio-only and ratio + family mixed-effects models.

**Supplemental Table S9.** Per-family regression of CK-TCS gene z-score on log (base/N-glucoside) ratio.

**Supplemental File 1.** Physiological measurements

**Supplemental File 2.** Metabolic data

**Supplemental File 3.** Gene counts and differentially expressed genes

**Supplemental File 4.** Total protein abundance and differentially enriched proteins

**Supplemental File 5.** Modeling, protoplasts and qRT-PCR experiments and primers

## Author contributions

O.H. and A.M.R. designed the experiments. M.C. performed and analyzed LC-MS proteomic data. I.P., M.S., and O.N. conducted cytokinin metabolic measurements. O.H. conducted physiological experiments and integration of the omics dataset. O.H. and A.M.R. wrote the manuscript, with input from co-authors.

## Data availability

All data are incorporated into this article and its online supplementary material. Customized R scripts are in the GitHub Repository (https://github.com/OmarHas07/RNA-ProteomeAnalaysis)

## Supplemental Figure Legends

**Supplemental Figure S1.**
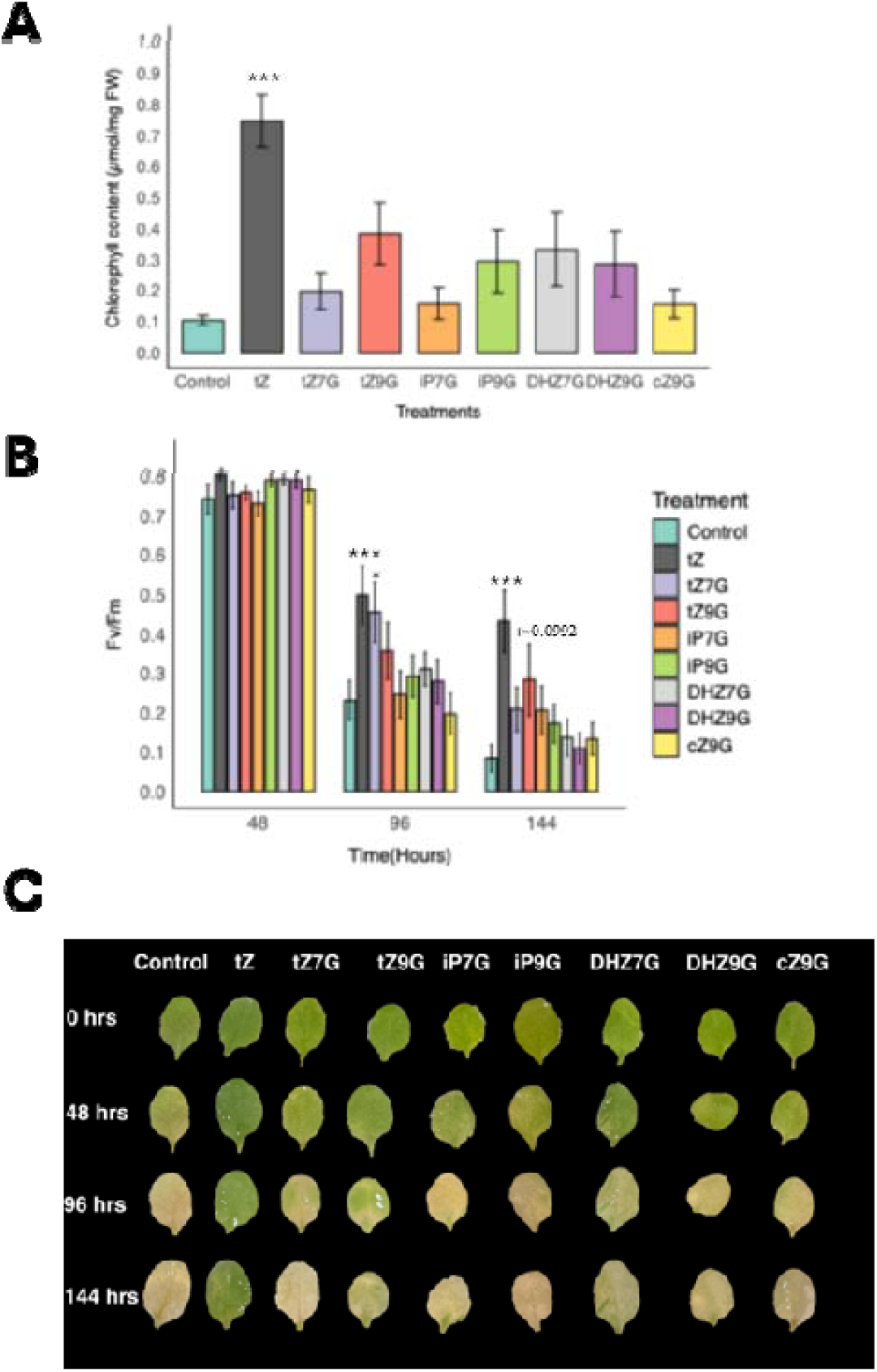
Activity of N-glucosides-treated leaves in dark-induced senescence assay on leaves 3-4. (**A-C**) Quantification of senescence progression across the 144-hour time course. Total chlorophyll content (µmol/mg FW) **(A),** Photosystem II efficiency (Fv/Fm) **(B)** of detached Arabidopsis leaves (positions 3-4) treated with 1 µM tZ, tZ7G, tZ9G, iP7G, iP9G, DHZ7G, DHZ9G, cZ9G, or negative control (NC) and incubated in the dark for 48, 96, and 144 hours. **(C)** Representative images of detached leaves from 4-week-old Arabidopsis plants treated with 1 µM of the indicated cytokinin bases form treatment across the time-course of dark-induced senescence (DIS). Data represent the mean ± SE of at least 4 biological replicates. Asterisks indicate statistical significance compared to the negative control (NC) as determined by one-way ANOVA followed by Bonferroni post-hoc test for (A-B) (*P < 0.05, **P< 0.001, ***P < 0.0001).

**Supplemental Figure S2.**
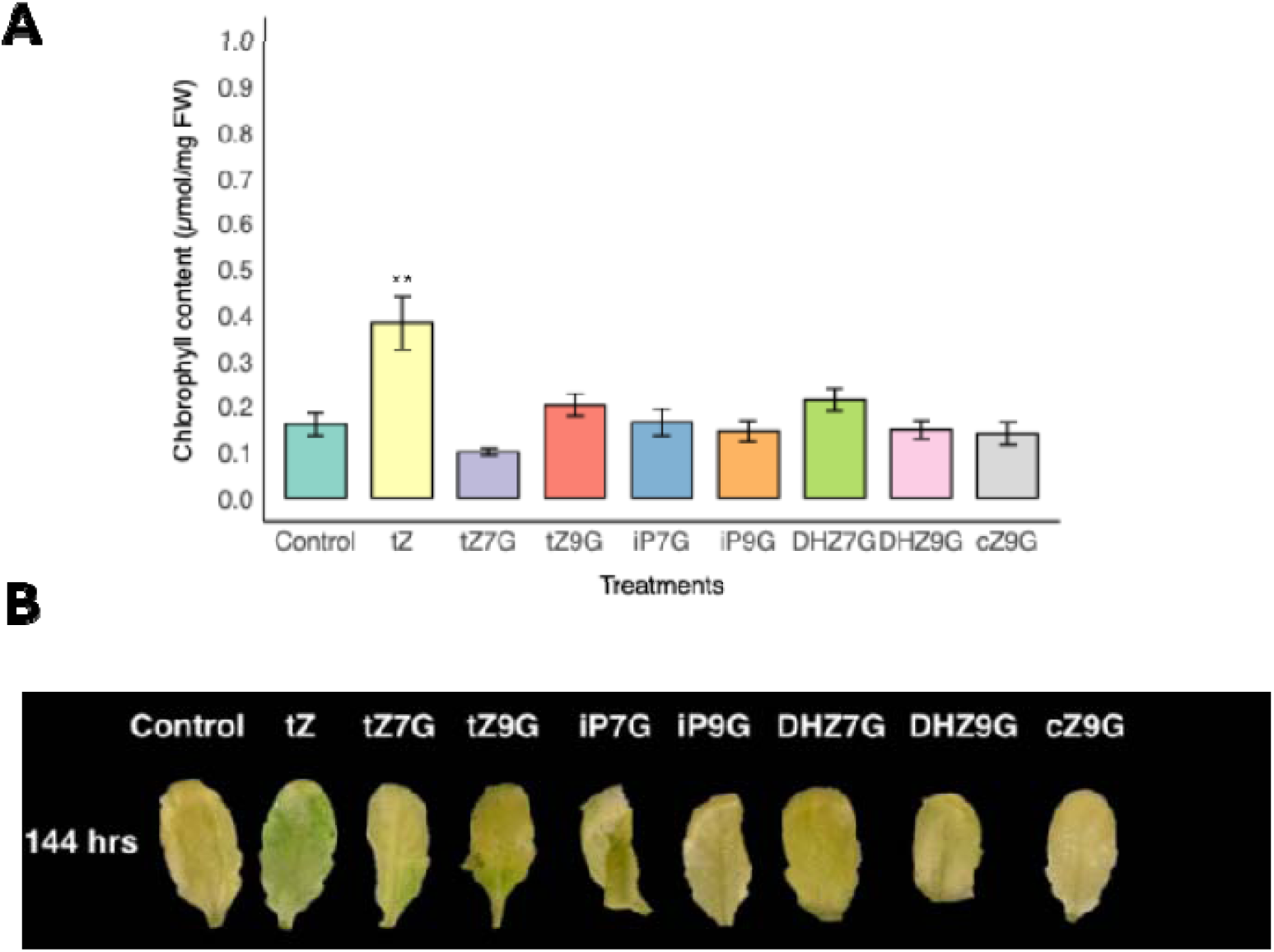
Activity of N-glucosides-treated leaves in dark-induced senescence assay on leaves 7-8. (**A-B**) Quantification of senescence progression across the 144-hour time course. Total chlorophyll content (µmol/mg FW) **(A)** of detached Arabidopsis leaves (positions 7-8) treated with 1 µM tZ, tZ7G, tZ9G, iP7G, iP9G, DHZ7G, DHZ9G, cZ9G, or negative control (NC) and incubated in the dark for 144 hours. **(B)** Representative images of detached leaves from 4-week-old Arabidopsis plants treated with 1 µM of the indicated cytokinin bases form treatment across the time-course of dark-induced senescence (DIS). Data represent the mean ± SE of at least 4 biological replicates. Asterisks indicate statistical significance compared to the negative control (NC) as determined by one-way ANOVA followed by Bonferroni post-hoc test for (*P < 0.05, **P< 0.001, ***P < 0.0001).

**Supplemental Figure S3.**
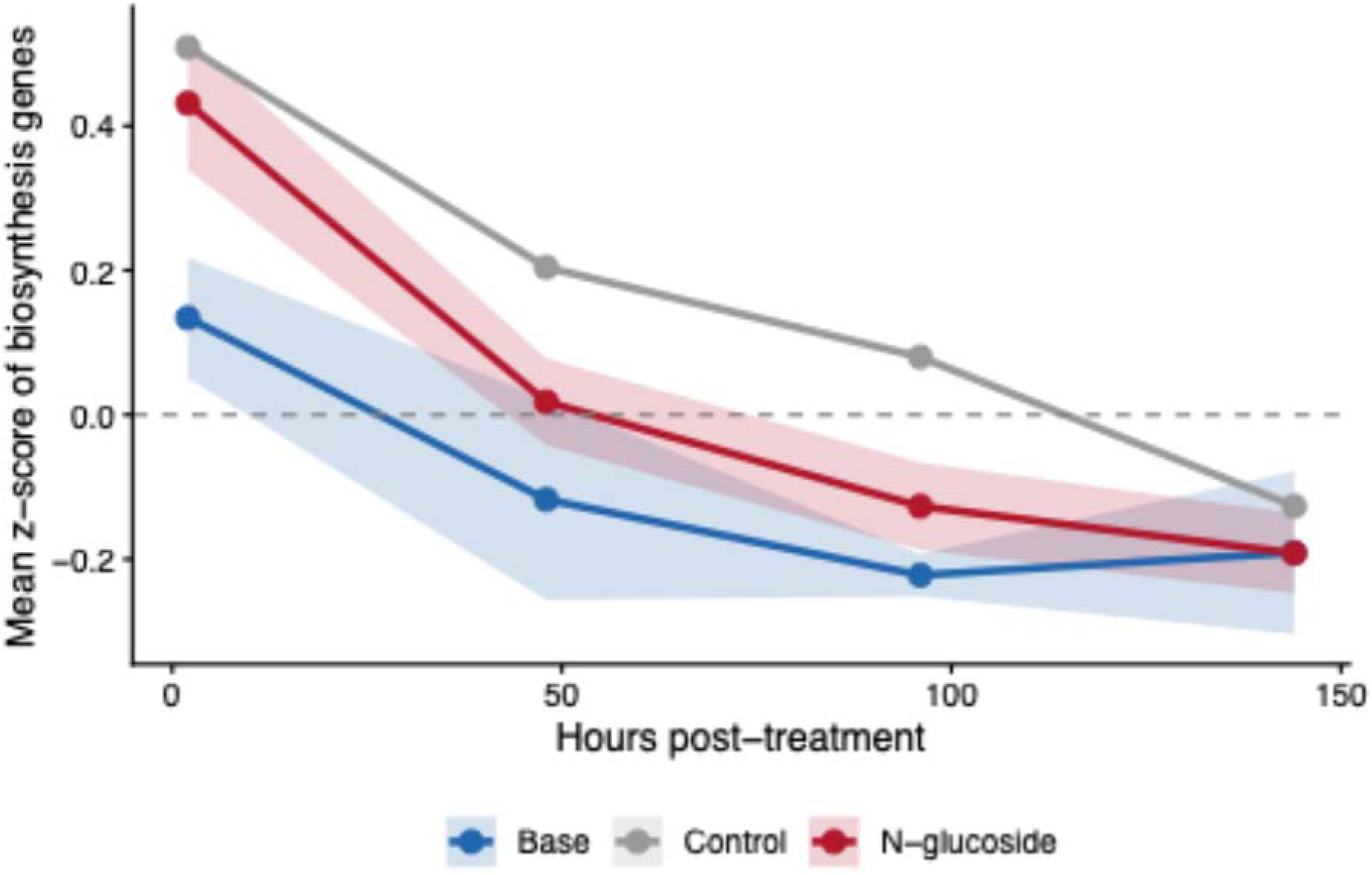
Average z-score of cytokinin biosynthesis-associated genes in response to N-glucoside and base form treatments. Line represents the average expression across time; ribbon represents the standard deviation across genes.

**Supplemental Figure S4.**
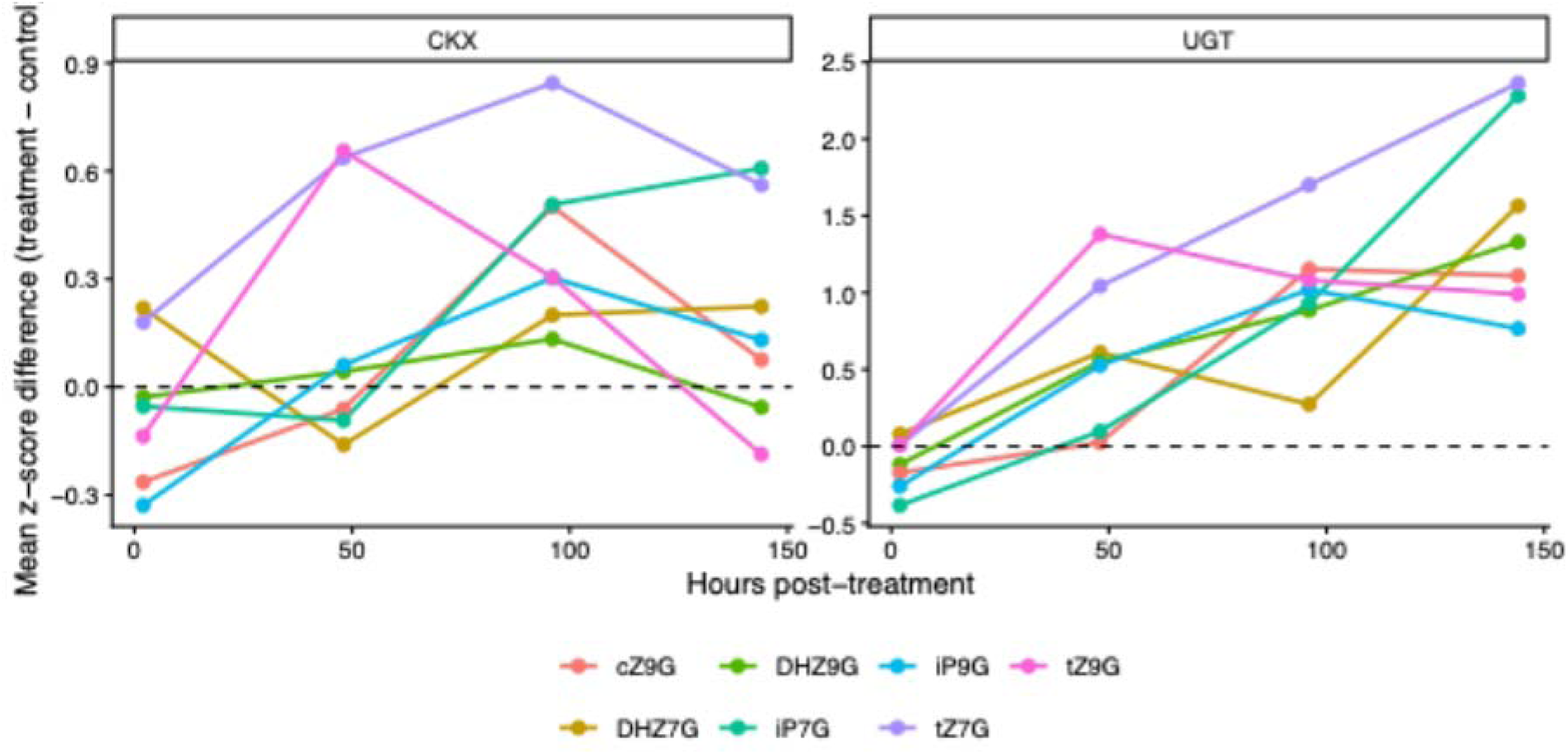
Expression of cytokinin degradation genes across time in response to N-glucoside and base form treatments. (Left) mean z-score expression of CKX genes while (Right) mean z-score expression of UGT genes across time points in response to N-glucosides and base forms.

**Supplemental Figure S5.**
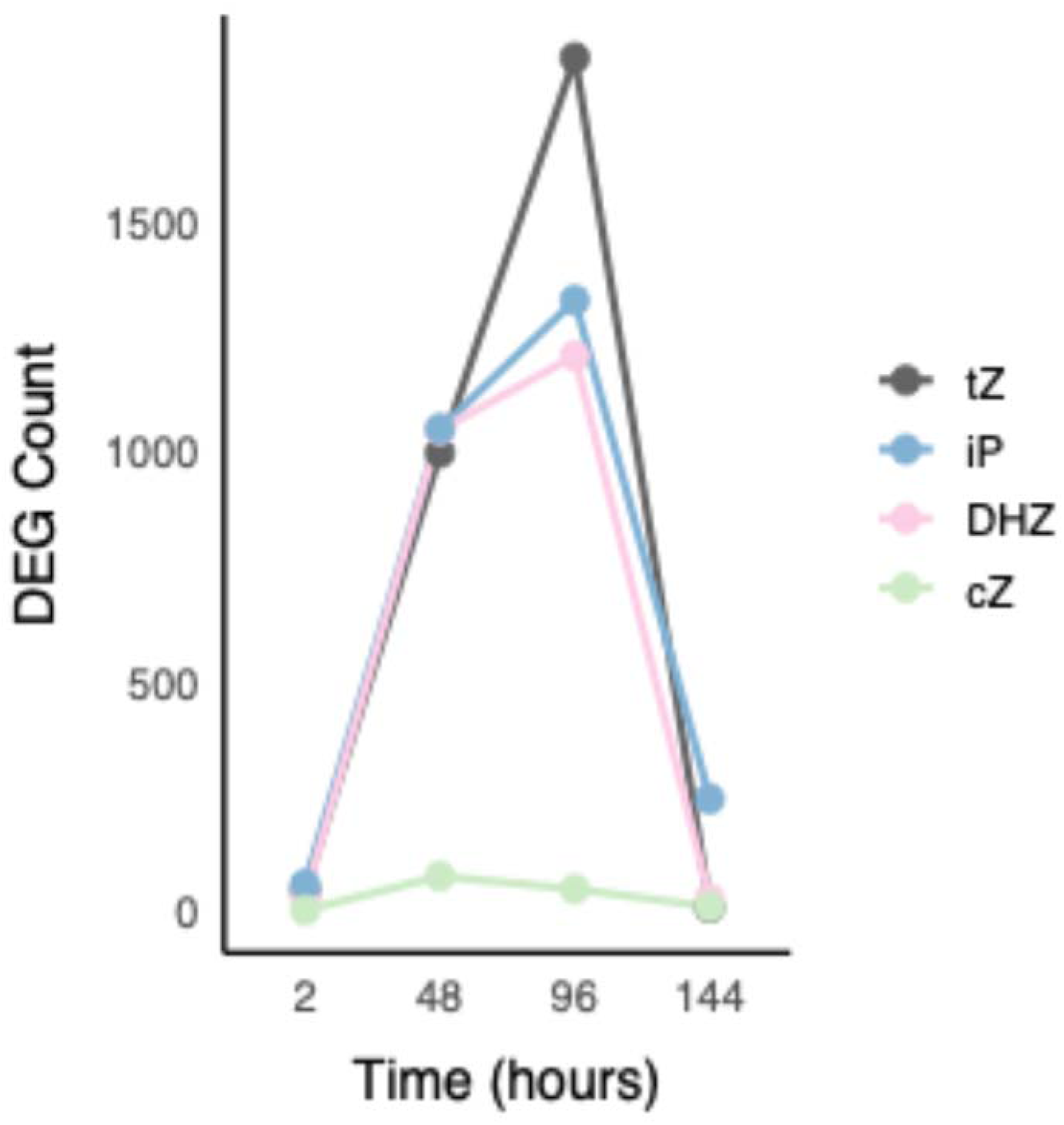
Number of Differentially Expressed Genes in response to base forms. FDR < 0.05 log2FC > 1

**Supplemental Figure S6.**
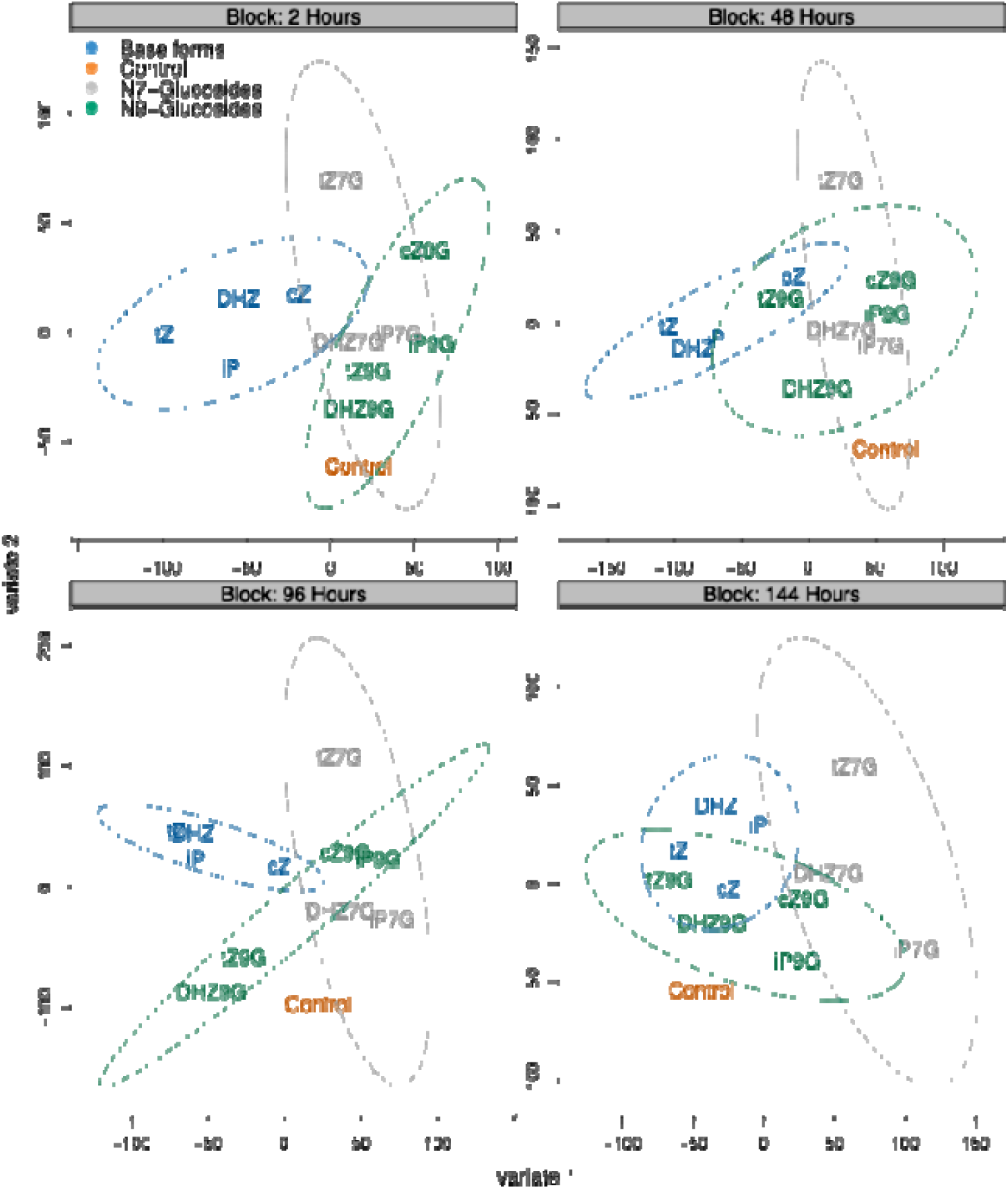
Block partial least square analysis comparing the global expression profiles of base form and N-glucoside treatments. Comparison of each individual time point showing that N-glucoside global expression pattern is substantially different compared to base forms.

**Supplemental Figure S7.**
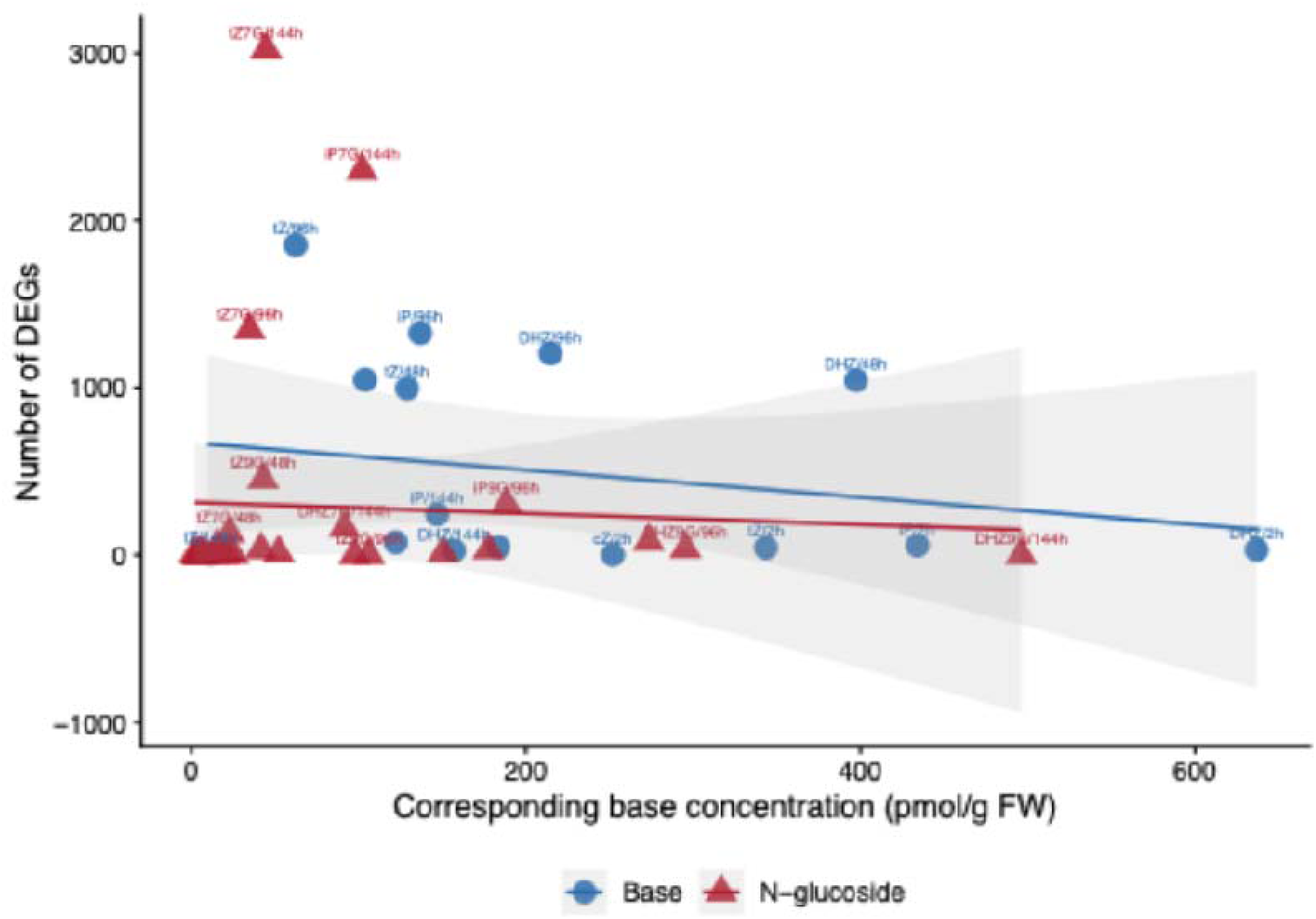
Plotting number of Differentially Expressed Genes versus accumulated base forms in response to exogenous base and N-glucoside treatments. DEGs in response to N-glucoside treatments are plotted against their corresponding base forms across different time points.

**Supplemental Figure S8.**
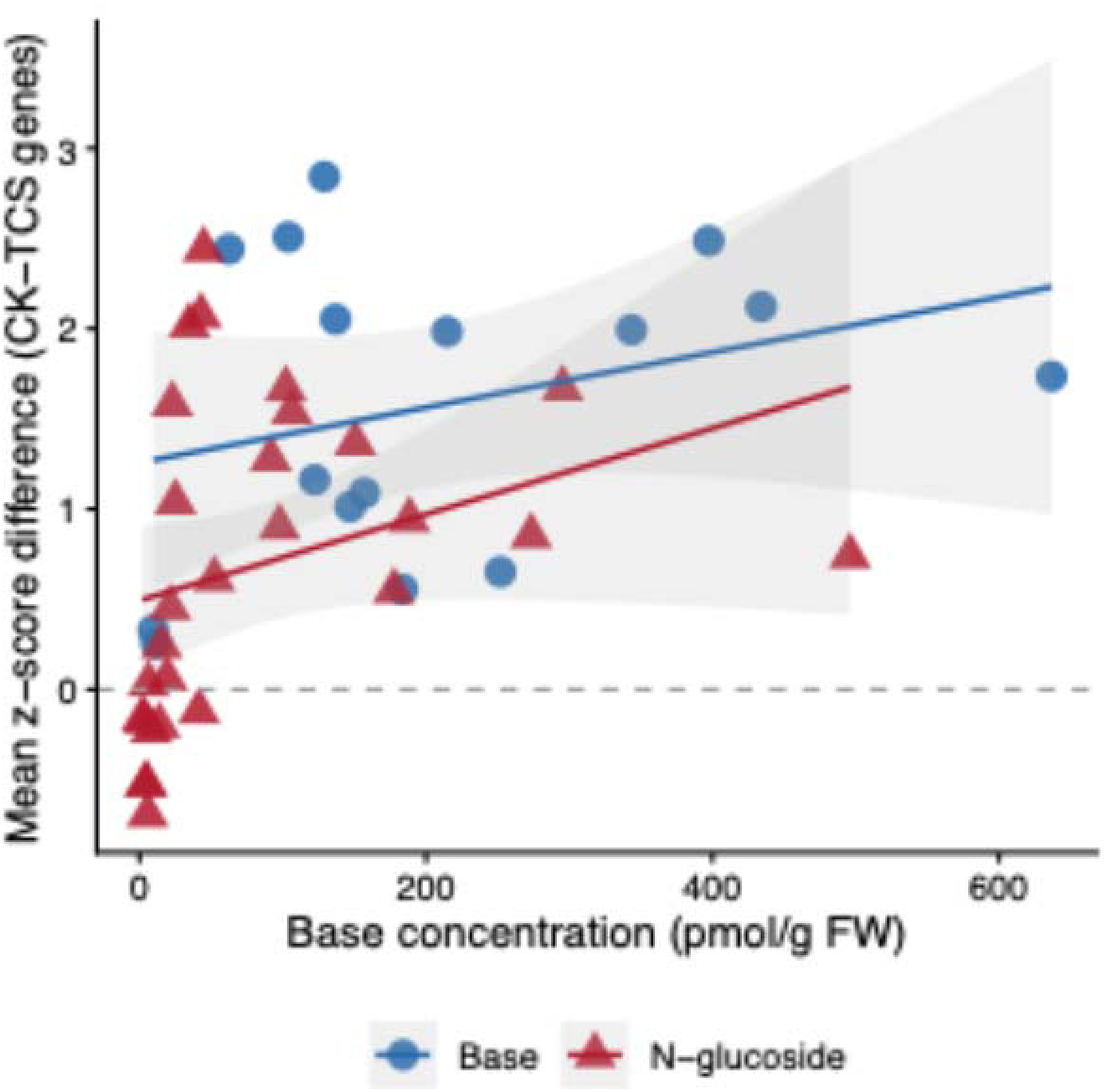
Plotting mean expression of Cytokinin Two Component Signaling Genes versus accumulated base forms in response to exogenous base and N-glucoside treatments.

**Supplemental Figure S9.**
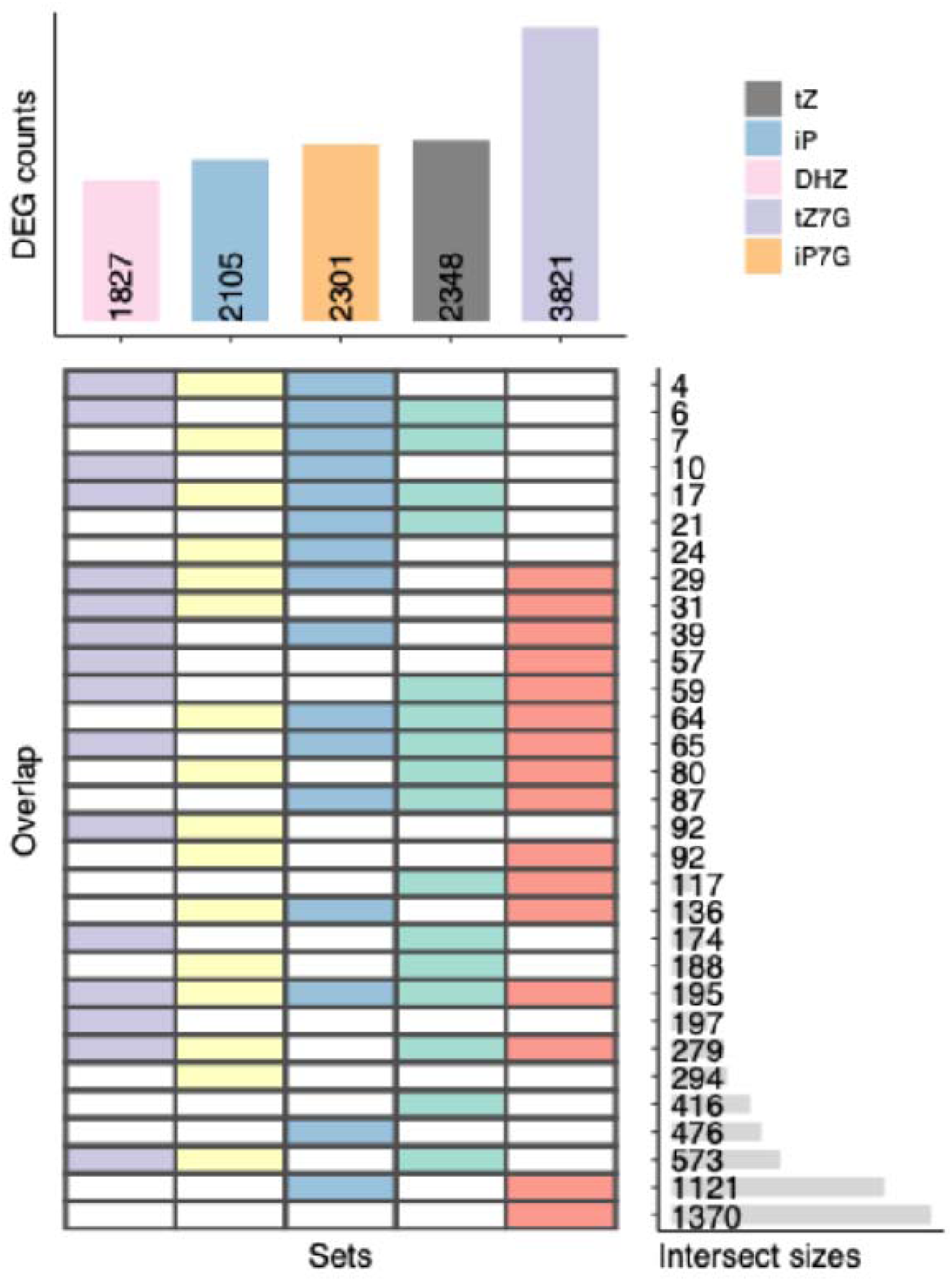
Upset plot showing the shared and unique numbers of Differentially Expressed Genes between tZ7G, iP7G and bioactive base forms.

**Supplemental Figure S10.**
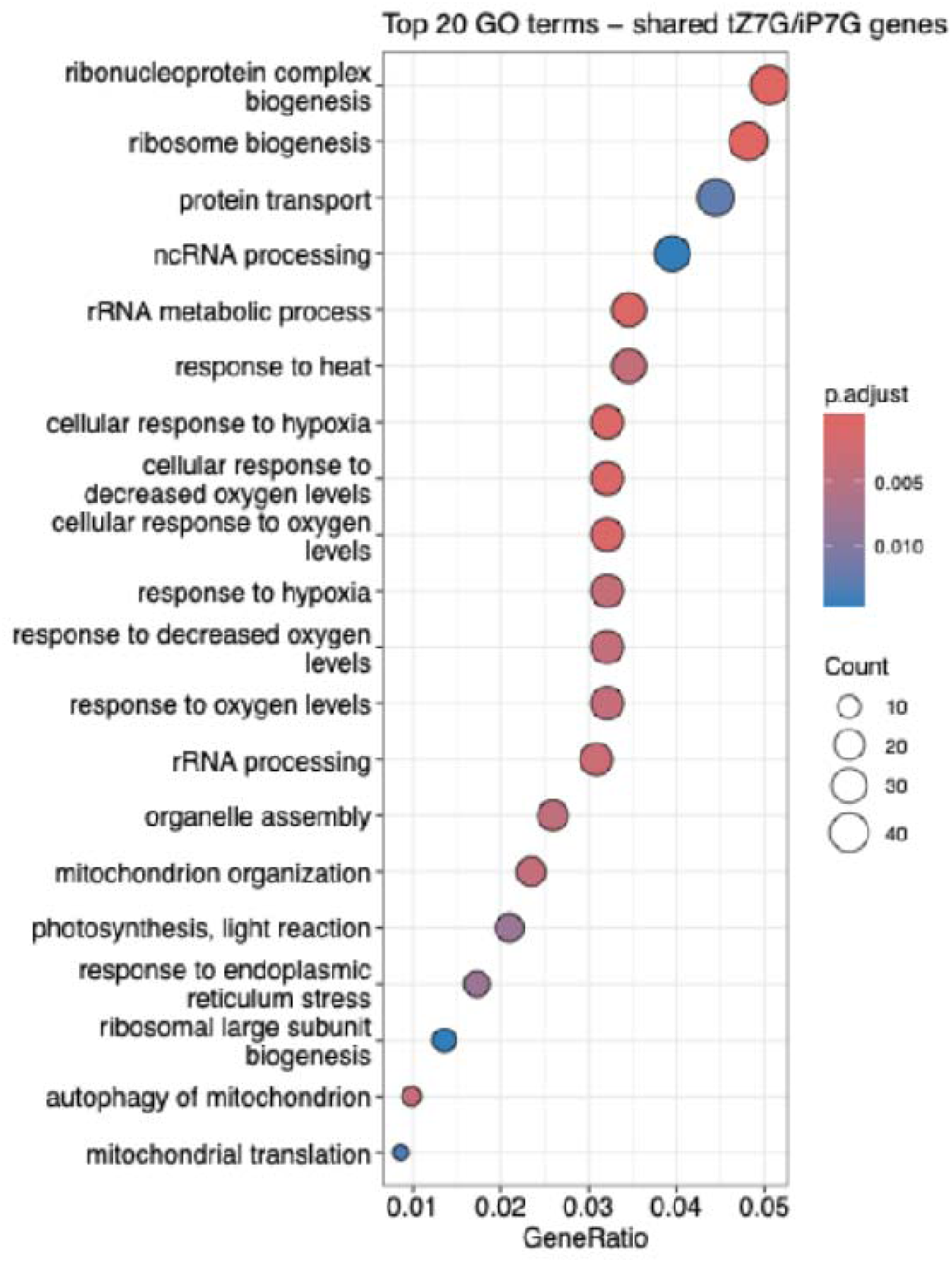
Gene Ontology analysis on unique genes co-regulated by tZ7G and iP7G but not by bioactive base forms.

**Supplemental Figure S11.**
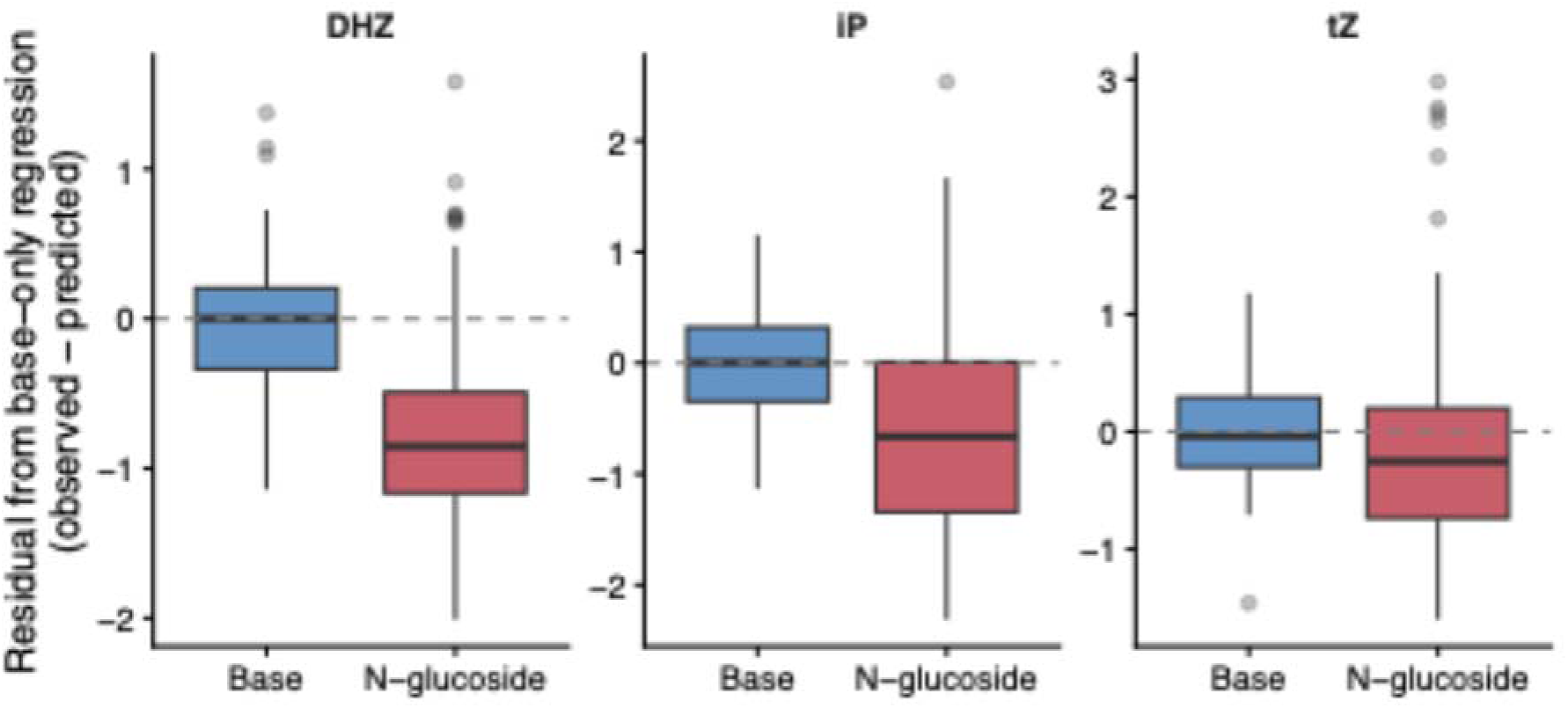
Residuals of linear regression modeling. Residuals from a base-only regression model applied to all treatment data within each family. N-glucoside-treated samples show significantly lower-than-predicted gene expression (Wilcoxon test against base residuals: tZ p = 0.047, iP p = 1.1 × 10□□, DHZ p = 1.2 × 10□□).

**Supplemental Figure S12.**
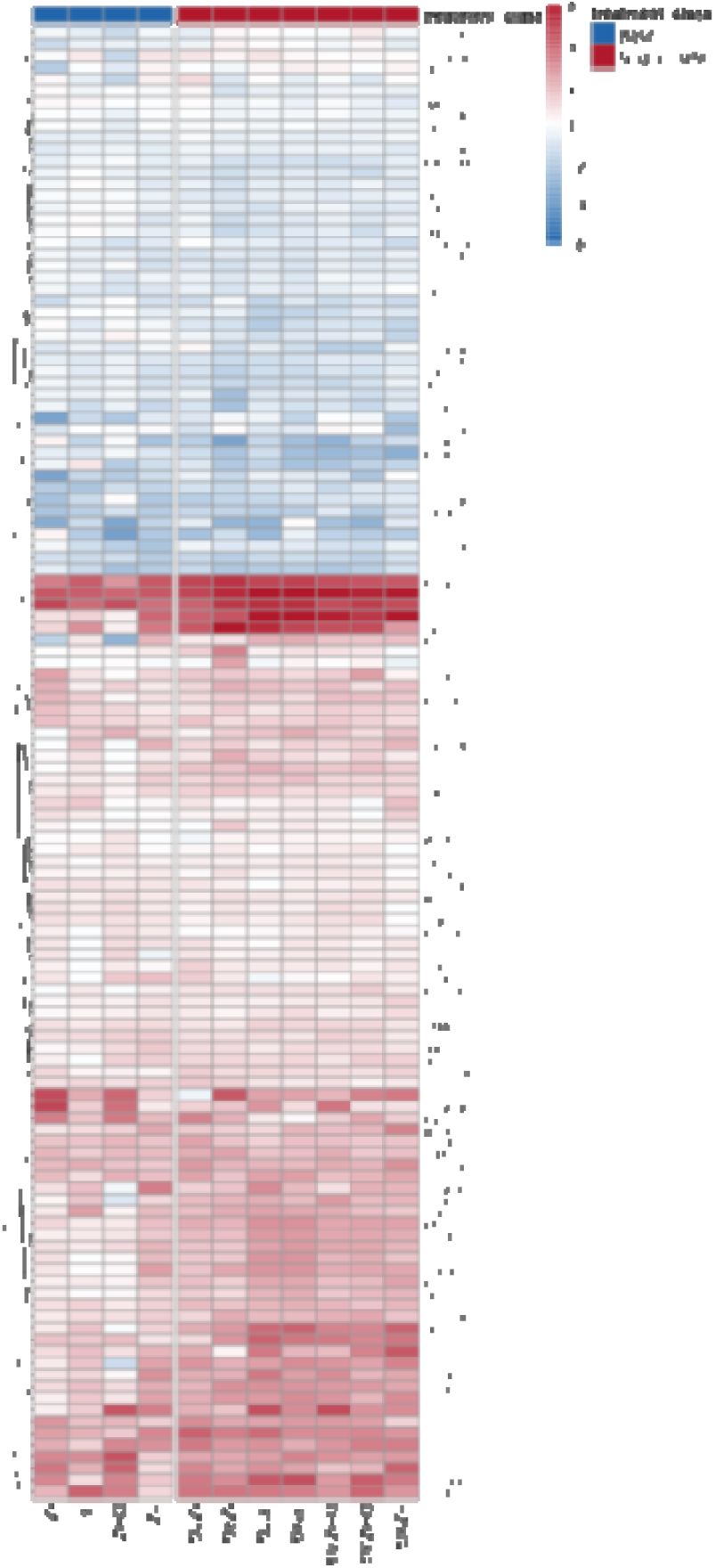
Heat map showing the differential protein abundance in N-glucosides and base form treated-samples. FDR < 0.05, Log2FC > 1.3

**Supplemental Figure S13.**
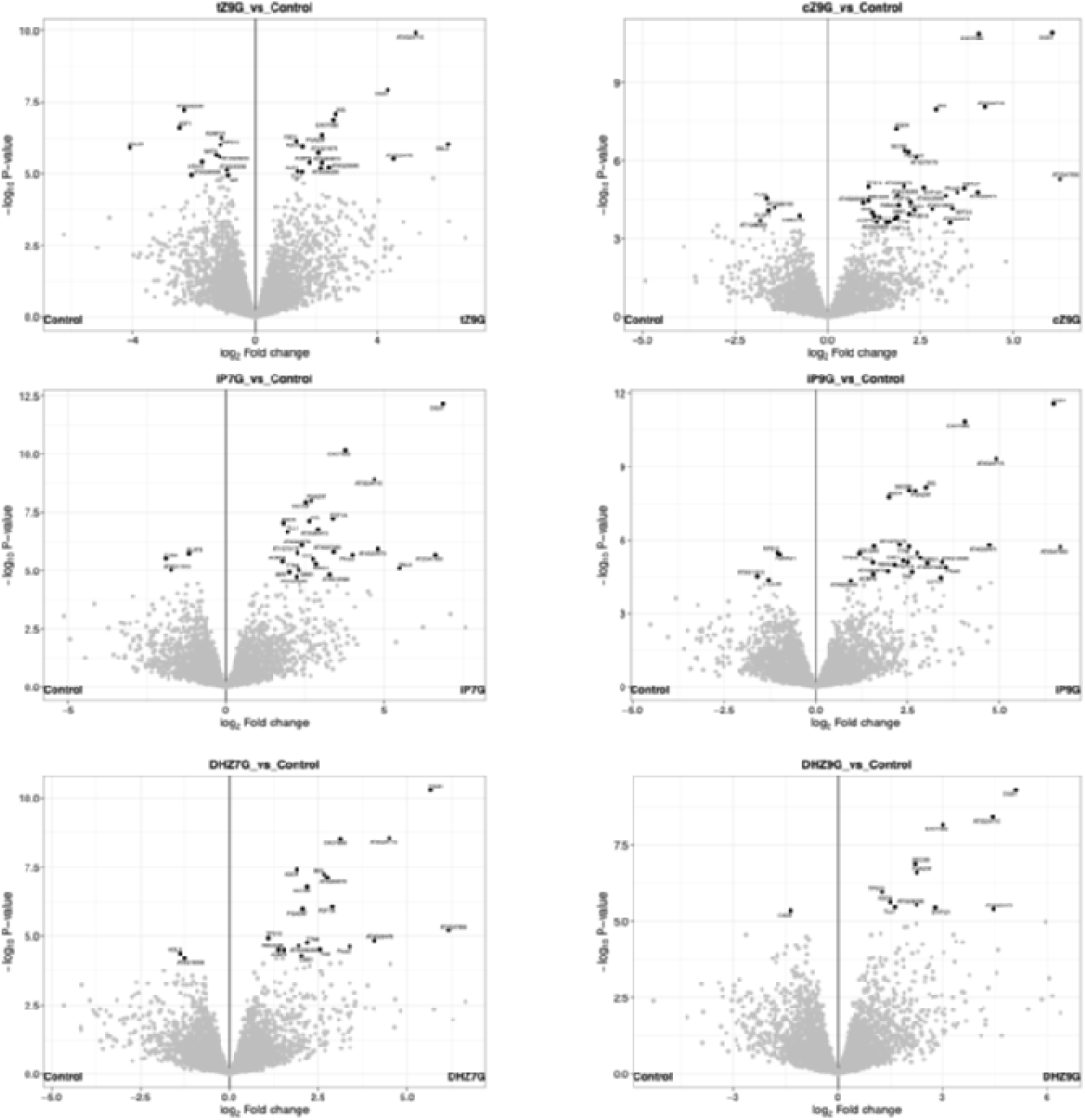
Volcano plots showing the differentially abundant proteins in response to N-glucoside-treated samples.

## Supplemental Tables

**Supplemental Table S1.** Tissue accumulation of N-glucoside conjugates across the time course. Concentration of each applied N-glucoside (pmol/g fresh weight) measured by LC-MS in detached Arabidopsis leaves treated with 1 µM of the indicated compound at 2, 48, 96, and 144 hours post-treatment. Values are mean ± SD across n = 5 replicates per treatment per timepoint (n = 4 for cZ9G at 2 h due to a missing sample). Letters denote groups not significantly different within a column based on one-way ANOVA followed by Tukey HSD pairwise comparisons (adjusted p < 0.05). Per-timepoint ANOVA results: 2 h, F□,□□ = 2.42, p = 0.053; 48 h, F□,□□ = 17.27, p = 3.0 × 10□□; 96 h, F□,□□ = 25.19, p = 4.5 × 10□¹□; 144 h, F□,□□ = 12.09, p = 1.1 × 10□□. Treatments sharing a letter within the same column are not significantly different from each other; treatments differing in letters are significantly different at adjusted p < 0.05.

| Treatment | 2 hours | 48 hours | 96 hours | 144 hours |
| --- | --- | --- | --- | --- |
| **tZ7G** | 212 ± 44 <u>ab</u> | 204 ± 68 <u>cde</u> | 275 ± 86 <u>ab</u> | 277 ± 102 <u>a</u> |
| **tZ9G** | 273 ± 162 <u>ab</u> | 397 ± 95 <u>ab</u> | 237 ± 165 <u>ab</u> | 338 ± 75 <u>ab</u> |
| **iP7G** | 125 ± 38 <u>a</u> | 71 ± 42 <u>c</u> | 126 ± 45 <u>a</u> | 141 ± 42 <u>a</u> |
| **iP9G** | 329 ± 123 <u>ab</u> | 345 ± 146 <u>ad</u> | 375 ± 84 <u>b</u> | 558 ± 142 <u>bc</u> |
| **DHZ7G** | 416 ± 259 <u>b</u> | 572 ± 126 <u>b</u> | 748 ± 86 <u>c</u> | 749 ± 278 <u>c</u> |
| **DHZ9G** | 268 ± 156 <u>ab</u> | 267 ± 60 <u>ade</u> | 224 ± 98 <u>ab</u> | 341 ± 103 <u>ab</u> |
| **cZ9G** | 150 ± 57 <u>ab</u> | 138 ± 47 <u>ce</u> | 145 ± 46 <u>a</u> | 213 ± 57 <u>a</u> |

**Supplemental Table S2.** Pool stability of N-glucosides across the 144-hour time course. Linear regression of N-glucoside pool concentration against time (h) for each of the seven N-glucoside treatments, fit on per-replicate measurements at 2, 48, 96, and 144 hours (n = 5 replicates per timepoint). Columns report the slope (pmol/g FW per hour), p-value of the slope coefficient (test against zero), regression R², percent change in pool concentration over the full time course (calculated as 100 × [final − initial] / initial using fitted regression endpoints), and a logical indicator of whether the slope was significantly negative (i.e., significant pool decline at p < 0.05). No N-glucoside treatment showed a statistically significant decline in pool concentration; iP9G (p = 0.011) and DHZ7G (p = 0.008) showed significant increases, indicating ongoing accumulation rather than depletion across the 144-hour window. The absence of pool decline despite measurable base release (Table S3, S4) indicates that N-glucoside levels are maintained by sources beyond passive consumption.

| Treatment | Slope pmol per hour | Slope pvalue | R squared | % change Over timecourse | significant decline |
| --- | --- | --- | --- | --- | --- |
| tZ7G | 0.55564143 | 0.09877577 | 0.1441026 | 30.1889653 | FALSE |
| tZ9G | 0.06925895 | 0.90721565 | 7.76E-04 | 23.8458715 | FALSE |
| iP7G | 0.22353956 | 0.27141983 | 0.06674618 | 13.0071638 | FALSE |
| iP9G | 1.52027441 | 0.01118949 | 0.30735883 | 69.6778469 | FALSE |
| DHZ7G | 2.47715353 | 0.0084844 | 0.32656853 | 80.2076319 | FALSE |
| DHZ9G | 0.37701475 | 0.4292811 | 0.03505429 | 27.3985105 | FALSE |
| cZ9G | 0.44191175 | 0.07706978 | 0.17242502 | 42.0599887 | FALSE |

**Supplemental Table S3.** Mass balance of N-glucoside conversion to free base over 144 hours. Quantitative comparison between excess free base accumulated in N-glucoside-treated leaves (compared with the time-matched Control treatment) and the corresponding N-glucoside pool concentration at the same timepoint. For each treatment at 144 hours, columns report: excess base (pmol/g FW), the cumulative mean concentration of free base above control attributable to N-glucoside conversion; % conversion needed (excess base ÷ initial N-glucoside pool × 100), the proportion of the initial N-glucoside pool that would need to have been converted to account for the observed excess base; % Gluc remaining (the fraction of the initial N-glucoside pool still present at 144 h, where 100% indicates a stable pool); and Gluc CVv (the coefficient of variation of the N-glucoside pool measurements at 144 h, expressed as a percentage). Values exceeding 100% in the “% conversion needed” column (DHZ9G: 186%, cZ9G: 188%) indicate that the cumulative base release exceeds the initial N-glucoside pool.

| Treatment | Timepoint | Excess base | % conversion needed | % Gluc remaining | Gluc CV |
| --- | --- | --- | --- | --- | --- |
| tZ7G | 144 | 43.4426 | 21.7946171 | 136.062113 | 37.018597 |
| tZ9G | 144 | 96.389 | 36.2225491 | 126.012111 | 22.204031 |
| iP7G | 144 | 100.185 | 85.2870569 | 117.361494 | 30.1566731 |
| iP9G | 144 | 175.701 | 53.7021574 | 169.88509 | 25.4337534 |
| DHZ7G | 144 | 90.0144 | 21.8106537 | 180.142881 | 37.0439513 |
| DHZ9G | 144 | 494.4586 | 186.462904 | 128.000136 | 30.0749339 |
| cZ9G | 144 | 269.4248 | 187.771382 | 145.769243 | 26.9614738 |

**Supplemental Table S4.** Conversion kinetics of N-glucosides to free bases under detached-leaf dark-induced senescence conditions. Apparent first-order conversion rate constants (k conv) calculated from the steady-state ratio of free base accumulation rate to mean N-glucoside pool concentration during the 2-144 h period. Columns report: G mean pmol, the time-averaged N-glucoside pool concentration (pmol/g FW); k conv/hour, the apparent first-order rate constant for conversion (% of pool per hour); half life hours, the calculated half-life under the assumption of first-order kinetics (h); and total converted 144h pct, the cumulative percentage of the initial pool that converted to base over the 144-hour time course. Treatments are ranked by k conv from fastest to slowest. DHZ9G and cZ9G show the fastest conversion rates (k conv ∼1.2-1.3% per hour), corresponding to half-lives of approximately 55-58 hours. tZ7G and DHZ7G show the slowest conversion (k conv ∼0.06-0.12% per hour), corresponding to half-lives of 568 and 1146 hours respectively. The 21-fold span of conversion rates across compounds indicates that N-glucoside-to-base conversion is highly compound-specific. All seven N-glucosides show measurable conversion across the time course, supporting the slow-release reservoir function described in the main text.

| Treatment | Base type | position | G mean pmol | % K conv/hour | Half life hours | Total converted 144h_pct |
| --- | --- | --- | --- | --- | --- | --- |
| DHZ9G | DHZ | 9G | 274.76765 | 1.26795897 | 54.6663729 | 182.586092 |
| cZ9G | cZ | 9G | 161.933526 | 1.19645996 | 57.9331699 | 172.290234 |
| iP7G | iP | 7G | 115.5105 | 0.57502665 | 120.54175 | 82.8038371 |
| iP9G | iP | 9G | 401.8805 | 0.35985153 | 192.620324 | 51.81862 |
| tZ9G | tZ | 9G | 311.25085 | 0.23342109 | 296.95139 | 33.6126374 |
| tZ7G | tZ | 7G | 242.01585 | 0.12213153 | 567.541537 | 17.5869408 |
| DHZ7G | DHZ | 7G | 621.15565 | 0.060504 | 1145.62212 | 8.71257568 |

**Supplemental Table S5.** Coefficients of the joint mixed-effects model relating standardized base concentration, standardized log□ (base/N-glucoside) ratio, and base family identity to CK-TCS gene z-score expression. Model formula: tcs_diff ∼ base_z + ratio_z + family + tp_factor + (1 | gene), fit by maximum likelihood (REML = FALSE) using lmer() from the lme4 package, with significance estimates from lmerTest. The dataset includes n = 440 observations spanning 11 treatments, 4 timepoints, and 10 canonical CK-responsive genes from the Bhargava et al. (2013) meta-analysis (≥13 datasets responding to CK). Predictors base_z and ratio_z are z-scored across the dataset (mean = 0, SD = 1), allowing direct comparison of standardized effect sizes. Family is a factor with levels (tZ, iP, DHZ, cZ), with tZ as the reference level (positive coefficients for other families indicate higher expression at the same predictor values; the negative familyiP, familyDHZ, and familycZ coefficients indicate that iP, DHZ, and cZ family treatments produce systematically lower TCS gene expression than tZ family at equivalent ratio values, consistent with the established potency hierarchy of cytokinin bases at AHK receptors). The standardized ratio coefficient (β = 0.45) is 2.6-fold larger than the standardized base coefficient (β = 0.17) and substantially more significant (t = 7.22, p = 2.4 × 10□¹² for ratio versus t = 2.64, p = 0.0087 for base), indicating that ratio is the dominant quantitative predictor of CK-TCS gene expression once family identity is accounted for.

| Term | Estimate | Std. Error | df | t value | Pr(>\|t\|) |
| --- | --- | --- | --- | --- | --- |
| (Intercept) | 1.12366969 | 0.1173186 | 35.75972 | 9.5779329 | 2.08E-11 |
| Base z | 0.17099686 | 0.06488 | 430 | 2.6355867 | 8.70E-03 |
| **Ratio z** | 0.45055107 | 0.06241651 | 430 | 7.2184603 | 2.40E-12 |
| **familyiP** | -0.629271 | 0.09155784 | 430 | -6.8729337 | 2.22E-11 |
| **familyDHZ** | -0.6204042 | 0.10950424 | 430 | -5.6655725 | 2.69E-08 |
| **familycZ** | -1.1516179 | 0.10251346 | 430 | -11.233822 | 7.50E-26 |
| **tp_factor48** | 0.66516711 | 0.10329337 | 430 | 6.4395915 | 3.21E-10 |
| **tp_factor96** | 0.45228016 | 0.1018836 | 430 | 4.4391849 | 1.15E-05 |
| **tp_factor144** | 0.04713445 | 0.10093681 | 430 | 0.4669699 | 6.41E-01 |

**Supplemental Table S6.** Comparison of mixed-effects models predicting CK-TCS gene z-score from base concentration, base-to-N-glucoside ratio, and base family identity. Seven candidate models compared via Akaike Information Criterion (AIC), Bayesian Information Criterion (BIC), log-likelihood, and marginal R². All models include time-point as a fixed factor and a random intercept for gene identity. Models tested: Family only (null), Base + Family, Base × Family (allowing per-family base slopes), Base + N-glucoside + Family, Ratio + Family, Ratio × Family (allowing per-family ratio slopes), and Base + Ratio + Family. Lower AIC indicates better fit; ΔAIC is calculated relative to the best-fitting model (Ratio × Family, AIC = 970). Direct head-to-head comparison of models with identical complexity but different quantitative predictors shows that Ratio + Family (AIC = 985) outperforms Base + Family (AIC = 1027) by ΔAIC = 42, and Ratio × Family (AIC = 970) outperforms Base × Family (AIC = 1012) by ΔAIC = 41, confirming ratio as the stronger quantitative predictor in both the parallel and per-family slope formulations. Marginal R² values rise from 0.16 (family only) to 0.49 (Ratio × Family).

| **Model** | **AIC** | **BIC** | **logLik** | **Marginal R2** | **Delta AIC** |
| --- | --- | --- | --- | --- | --- |
| **Ratio x Family** | 970 | 1023 | -472 | 0.489 | 0 |
| **Base + Ratio + Family** | 980 | 1025 | -479 | 0.475 | 9.68 |
| **Ratio + Family** | 985 | 1026 | -482 | 0.467 | 14.6 |
| **Base x Family** | 1012 | 1065 | -493 | 0.444 | 41.4 |
| **Base + Glucoside + Family** | 1023 | 1068 | -501 | 0.426 | 52.9 |
| **Base + Family** | 1027 | 1068 | -503 | 0.419 | 56.9 |
| **Family only (null)** | 1199 | 1235 | -590 | 0.164 | 229 |

**Supplemental Table S7.** Comparison of ratio-only versus ratio + family versus ratio × family mixed-effects models. Three nested mixed-effects models compared on the same dataset. The Ratio only model includes only the standardized log□(base/N-glucoside) ratio and timepoint as fixed effects, with random intercept for gene. The Ratio + Family model adds family as a fixed effect (parallel slopes assumption: families share a common ratio➔response slope but may have different intercepts). The Ratio × Family model further adds a ratio × family interaction (per-family ratio➔response slopes). Adding family identity to a ratio-only model substantially improves fit (ΔAIC = 104) and adding family-specific slopes provides further improvement (ΔAIC = 15). The progressive improvement indicates that while ratio captures the dominant quantitative relationship, base family identity contributes additional explanatory power consistent with intrinsic differences in cytokinin potency at AHK receptors. The Ratio × Family model is the best-fitting overall (lowest AIC).

| Model | AIC | BIC | logLik | delta_AIC |
| --- | --- | --- | --- | --- |
| **Ratio only** | 1089 | 1117 | -537 | 119 |
| **Ratio + Family** | 985 | 1026 | -482 | 14.6 |
| **Ratio \* Family (interaction)** | 970 | 1023 | -472 | 0 |

**Supplemental Table S8.** Likelihood ratio test comparing ratio-only and ratio + family mixed-effects models. Likelihood ratio test (LRT) of nested mixed-effects models predicting CK-TCS gene z-score, performed via anova(m_ratio, m_ratio_fam). Columns report number of parameters (npar), Akaike Information Criterion (AIC), Bayesian Information Criterion (BIC), log-likelihood (logLik), deviance (–2 × log-likelihood), chi-squared statistic of the LRT, degrees of freedom of the test, and the p-value. The reduced model (m_ratio) includes log□(base/N-glucoside) ratio and timepoint as fixed effects with random intercept for gene; the full model (m_ratio_fam) adds family as a fixed factor with three additional parameters (one per non-reference family level).

| Model | npar | AIC | BIC | logLik | -2*log(L) | Chisq | Df | Pr(>Chisq) |
| --- | --- | --- | --- | --- | --- | --- | --- | --- |
| **m_ratio** | 7 | 1088.7 | 1117.3 | -537.37 | 1074.7 |  |  |  |
| **m_ratio_fam** | 10 | 984.7 | 1025.6 | -482.35 | 964.7 | 110.03 | 3 | < 2.2e-16 *** |

**Supplemental Table S9.** Per-family regression of CK-TCS gene z-score on log□ (base/N-glucoside) ratio. Linear regression of standardized z-score CK-TCS gene expression on log (base/N-glucoside) ratio, fit separately within each base family (tZ, iP, DHZ, cZ) and including timepoint as a covariate. Columns report number of observations within the family (n_obs), the slope coefficient on log□_ratio with standard error and p-value, and the regression R². iP and DHZ families show nearly identical ratio-response slopes (0.32 and 0.33 respectively) and similar R² (0.46-0.52). The tZ family shows a shallower slope (0.22) but high R² (0.50), indicating that within tZ samples, ratio remains a strong predictor but the dynamic range of response per unit log□ ratio change is smaller. The cZ family shows the shallowest slope (0.16) and lowest R² (0.33).

| family | N obs | slope | Slope se | Slope p | R squared |
| --- | --- | --- | --- | --- | --- |
| **tZ** | 120 | 0.224 | 0.0311 | 0 | 0.495 |
| **iP** | 120 | 0.324 | 0.0303 | 0 | 0.518 |
| **DHZ** | 120 | 0.326 | 0.0342 | 0 | 0.458 |
| **cZ** | 80 | 0.159 | 0.0311 | 0 | 0.326 |

